# Rapid degradation of 6 class I HDAC complexes reveals minimal functional overlap between complexes

**DOI:** 10.1101/2025.08.21.671480

**Authors:** Robert E. Turnbull, Oksana Gonchar, David M. English, Tom A. Wright, India M. Baker, Kristupas Sirvydis, Shaun M. Cowley, John W.R. Schwabe

## Abstract

The class 1 HDACs 1, 2 and 3 form seven families of distinct large multiprotein complexes that regulate gene expression via deacetylation of lysines in histone tails. The degree of redundancy and functional overlap between complexes and their primary gene targets, remains unknown. We used CRISPR/Cas9 to independently tag HDAC complexes with FKBP12^F36V^ in HCT116 cells enabling rapid (<1 hr), PROTAC-mediated, degradation. RNA sequencing at 6 h reveals that together, the 4 major complexes (CoREST, NuRD, NCoR/SMRT and SIN3A) perturbed >50% of expressed genes. More than 60% of these are specific to an individual complex. Of genes regulated by more than one complex, approaching 50% are reciprocally regulated such that HDAC complexes act as antagonistic regulators. Homer analysis strongly suggests that the complexes are reliant on different transcription factors. This is the first study to identify the primary targets of individual HDAC complexes and directly compare the effects of rapid degradation on gene regulation in the same biological system.

## Introduction

Class 1 histone deacetylases (HDACs) are zinc-dependent enzymes that catalyse the removal of acetyl groups from the side chain of lysine residues, predominantly within histone tails. HDACs 1 and 2 have 80% sequence identity and are largely interchangeable. They form the catalytic component of six multi-protein complexes: CoREST, MiDAC, MIER, NuRD, Sin3 and RERE (Bantscheff *et al*, 2011; Ding *et al*, 2003; Hassig *et al*, 1997; You *et al*, 2001; Zhang *et al*, 1998; Zoltewicz *et al*, 2004). The other major nuclear HDAC, HDAC3, forms the catalytic subunit of the NCoR/SMRT complex (figure 1A) (Guenther *et al*, 2000).

**Figure 1.**
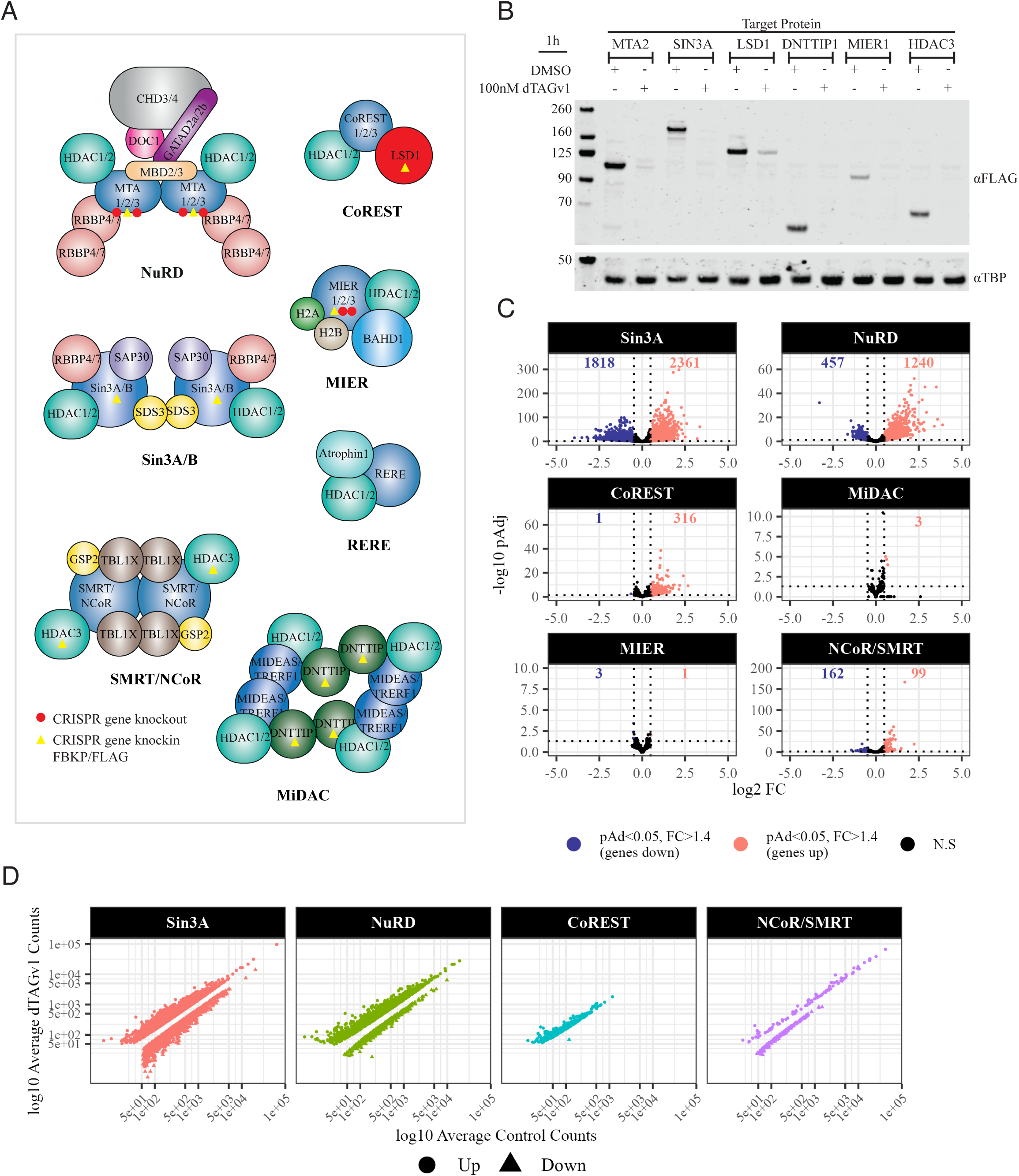
Rapid, PROTAC-mediated degradation of targeted HDAC complexes results in gene expression changes after 6h. A Schematic of the 7 class 1 HDAC complex families. Unique proteins targeted for CRISPR/Cas9 are identified with a yellow triangle and paralogues that were knocked out identified with a red circle. B Western blot for the FLAG epitope, identifying the six targeted proteins in six HCT116 cell lines in the presence or absence of dTAG^V^-1 (100 nM). An equal volume of DMSO was used for controls. TATA binding protein was used as a loading control. C Volcano plots for Log_2_ fold change (log_2_ FC) and log_10_ adjusted p-value (-log_10_ pAdj) calculated from the RNA sequencing analysis. Significant genes were classified as having an pAdj <0.05 and log_2_ FC ≥0.485 (FC 1.4). RNA sequencing was performed using biological triplicates and significance calculated using the DESEQ2 package with APEGLM shrinkage. D Average normalised counts for control and dTAG^V^-1 triplicates of significant genes for each complex.

These complexes play essential roles during embryogenesis as knockout of unique proteins exhibit embryonic (CoREST, MiDAC, NCoR/SMRT, RERE, SIN3A, NuRD) (Bhaskara *et al*, 2008; Dannenberg *et al*, 2005; Hendrich *et al*, 2001; Lu *et al*, 2008; Turnbull *et al*, 2020; Wang *et al*, 2007; Zoltewicz *et al*., 2004) or perinatal (MIER) (Lakisic *et al*, 2016) lethality at various stages of development. Since all complexes are required for normal development, this suggests a lack of redundancy. However, in cell models of differentiation, the effects are less pronounced, with the loss of individual complexes often producing similar phenotypes. For instance, depletion of either CoREST, NuRD or NCoR/SMRT in mouse embryonic stem cells (mESCs) causes disruptions in embryoid body formation and abnormal differentiation patterns (Burgold *et al*, 2019; Foster *et al*, 2010; Malla *et al*, 2024; Wang *et al*, 2009a). Similarly, knockdown of CoREST in mESCs results in decreased expression of cardiac lineage markers (Foster *et al*., 2010) and depletion of MiDAC in human induced pluripotent stem cells (hiPSCs) prevents their differentiation into mature cardiomyocytes (Lu *et al*, 2024).

In somatic and disease models there is further evidence of overlapping functions of the different complexes. For example, cooperation between CoREST and SIN3 suppresses neuronal gene expression in non-neuronal cells (Andres *et al*, 1999; Huang *et al*, 1999; Naruse *et al*, 1999). Interaction between CoREST and NuRD has been shown to regulate inner ear development (Patel *et al*, 2018). In cancer, mesenchymal transcription factors recruit several HDAC complexes to repress epithelial gene expression, thereby promoting epithelial-mesenchymal transition (EMT) (Lin *et al*, 2010a; Peinado *et al*, 2004). CoREST and MiDAC have also been shown to interact and promote EMT in non-small cell lung cancer (Liu *et al*, 2022). Interestingly, these complexes can also have antagonistic effects; for instance, SIN3A inhibits, while SIN3B promotes breast cancer cell proliferation and metastasis (Lewis *et al*, 2016). Taken together, these studies suggest that class 1 HDAC complexes in some circumstances have independent roles but also that inter-complex relationships occur, and these may act in a cooperative or antagonistic manner.

To date, understanding the diverse roles of these complexes has been limited due to studies focusing on individual complexes in diverse cell systems. Investigations into HDAC complex function have primarily relied on knockdown, knockout or inhibitor-based approaches (Millard *et al*, 2017). While knockdown and knockout approaches are specific, they only reveal the medium- to long-term effects of complex loss. In contrast, inhibitor approaches are faster but are limited by their lack of complex specificity. To address these limitations, we used CRISPR/Cas9 to incorporate the PROTAC target FK506-binding protein 12 (FKBP12^F36V^) and FLAG epitope onto endogenous alleles of unique HDAC complex components in HCT116 cells (Boija *et al*, 2018; English *et al*, 2024b; Jinek *et al*, 2012; Liu *et al*, 2021; Nabet *et al*, 2020; Ran *et al*, 2013; Weintraub *et al*, 2017).

We successfully independently targeted six class 1 HDAC complexes in HCT116 cells, achieving degradation of the targeted complex proteins within 1h of PROTAC treatment. RNA sequencing at 6 hours post-treatment revealed a dynamic landscape of differentially expressed genes (DEGs), with over 50% of all genes in HCT116 cells perturbed by the four main complexes: CoREST, NCoR/SMRT, NuRD and Sin3A. Notably, more than 60% of DEGs were specific to individual complexes. Overlapping genes whose expression changed in the same direction were largely restricted to pairs of complexes. Interestingly, we observed a substantial overlap of genes whose expression changed in opposite directions between several complexes. Pathway analysis of these gene sets revealed enrichment for neuronal pathways that suggest, SIN3A promotes, while CoREST and NuRD repress, gene expression. HOMER analysis identified several sets of genes enriched for transcription factor (TF) binding; however, these showed minimal overlap between the complexes and, apart from SIN3A, were not associated with changes in TF expression. This study represents the first comparative analysis of complex-specific early gene targets for all 6 HDAC1/2 complexes and highlights a reciprocal relationship in gene regulation.

## Results

### FKBP12^F36V^-tagged HDAC complexes are expressed at wild-type levels and retain enzymatic activity

An FKBP12^F36V^ degradation tag (dTAG) and a FLAG epitope were added to the endogenous genes for specific subunits of six class 1 HDAC complexes independently in HCT116 cells (see methods, Figure 1A). For NuRD and MIER complex targeting, MTA1 and 3, and MIER2 and 3, respectively were knocked out to eliminate potential redundancy. The expression levels of targeted SIN3A, LSD1, DNTTIP1 and HDAC3 proteins were similar to those observed in parental HCT116 cells (Figure S1A). Expression of MIER1 and MTA2 targeted proteins was reduced but is in line with earlier studies (Figure S1A) (Eaton *et al*, 2018). Proliferation rates of the targeted HCT116 cells were comparable to those of parental cells with the exception of the tagged NuRD cell line which was reduced (Figure S1B). Since the NuRD complex is known to be involved in cell proliferation, the observed reduction likely reflects decreased levels of overall NuRD complex in the absence of MTA1 and MTA3 (Jiang *et al*, 2025; Kumar *et al*, 2011). We confirmed that the targeted proteins assemble into complexes with the expected HDAC activity (Figure S1C) and, for CoREST, demethylase activity (Figure S1D). These results indicate that introducing the FKBP12^F36V^/FLAG tags (∼13 kDA) into endogenous alleles does not significantly impair the physiological expression or function of these complexes.

### Targeted HDAC1/2 complex components are rapidly degraded upon addition of dTAG^V^-1

The FLAG epitope was readily detected in all lines expressing the CRISPR-modified endogenous protein (Figure 1B). The relative abundance of targeted proteins is consistent with their RNA expression levels (Figure S2A) and similar to findings from previous studies (Alshehri *et al*, 2025; Vcelkova *et al*, 2023). After just 1 h incubation following dTAG^V^-1 treatment, we observed >95% reduction in all target proteins (Figure 1B), demonstrating the effectiveness of this system for rapid endogenous protein degradation. Notably, the speed of degradation observed is quicker than that reported for other endogenously targeted proteins (Liu *et al*., 2021) but is comparable with that observed for stably expressed HDAC1-FKBP12^F36V^/FLAG (English *et al*, 2024a).

LSD1 and RCOR1/2/3, and DNTTIP1 and MIDEAS form components of the CoREST and MiDAC complexes, respectively. Previous studies have shown that the stability of these proteins depends on their incorporation into the respective complexes (Foster *et al*., 2010; Mondal *et al*, 2020; Turnbull *et al*., 2020). Upon treatment with dTAG^V^-1 for 6 h, we observed a ∼50% reduction in RCOR1 and MIDEAS levels in the corresponding cell line, with no further decrease detected after 24 h (Figure S1E and F), demonstrating complex partner co-dependency in these cells.

### Disruption of individual HDAC complexes reveals that the magnitude and direction of gene expression changes is highly complex dependent

RNA sequencing following 6 h of dTAG^V^-1 treatment revealed substantial, complex-dependent variation in the number of differentially expressed genes (DEGs) (pAdj <0.05, log2FC >=0.485 (FC>=1.4)). Depletion of the SIN3A and NuRD complexes perturbed the expression of thousands of genes (4179 and 1697 DEGs respectively). Depletion of CoREST and NCoR/SMRT resulted in fewer gene expression changes (317 and 261 DEGs respectively). Depletion of the MiDAC and MIER complexes resulted in very few gene expression changes (3 and 4 DEGs respectively) (Figure 1C).

In apparent contrast to these findings in HCT116 cells, previous studies have reported substantial gene expression changes in MiDAC knockout cells (Mondal *et al*., 2020; Turnbull *et al*., 2020) and a role for MIER1 in regulating gene expression during mouse liver regeneration (Chen *et al*, 2023). The fact that very few genes are perturbed in HCT116 cells suggests either that these complexes may not play a role in this terminally differentiated cancer cell line or that previous observations reflect secondary effects due to long-term loss or perhaps that these complexes play a more important role in differentiating cells during development (Turnbull *et al*., 2020; Wang *et al*, 2023). Given the limited effect of MiDAC and MIER1 depletion in this cell line, we did not include these data in further analyses.

NuRD and CoREST resulted in more upregulated than downregulated DEGs (Figure 1C). The ratio of up- to downregulated genes, however, differed between complexes. Only 1% of DEGs were downregulated for CoREST and 27% for NuRD. Strikingly, SIN3A differed from NuRD and CoREST complexes in that there was a near-equal distribution of upregulated (2,361) and downregulated (1,818) genes (Figure 1C), suggesting the SIN3A complex plays dual roles in both gene activation and repression. This is consistent with a broad function in transcriptional regulation (Adams *et al*, 2018; Mahendrawada *et al*, 2025; Tiana *et al*, 2018). Interestingly, 62% of DEGs following NCoR/SMRT depletion were downregulated (Figure 1C), despite NCoR/SMRT being a co-repressor typically linked to gene silencing. A similar pattern was previously observed in a Drosophila HDAC3 knockout embryo model (Tang *et al*, 2023).

Normalised mean control versus dTAG^V^-1 counts of DEGs showed complex specific distribution. SIN3A and NuRD depletion affected genes irrespective of control expression levels or whether they were up- or downregulated (Figure 1D). In contrast, CoREST depletion predominantly led to upregulation of genes with below 500 counts in control cells. Interestingly, NCoR/SMRT depletion significantly increased expression of genes with control counts greater than 500, while repression following NCoR/SMRT depletion was restricted to genes with expression below 500 counts (Figure 1D). Again, similar findings for NCoR/SMRT were observed by Tang et al. (Tang *et al*., 2023).

### Each HDAC complex regulates a specific set of genes with limited overlap

After filtering out genes expressed with lower than 50 counts, ∼12,000 genes are expressed in the HCT116 cell line. Of these, 6,461 were significantly perturbed following degradation of one or other HDAC complexes. Most of these genes (5281) are uniquely perturbed by a single complex. For SIN3A, 85% of upregulated genes and 92% of downregulated genes are unique to this complex (Figure 2A). For NuRD, CoREST and NCoR/SMRT approximately two-thirds of the upregulated genes and ∼75% of downregulated genes are uniquely regulated by each complex (Figure 2A).

**Figure 2.**
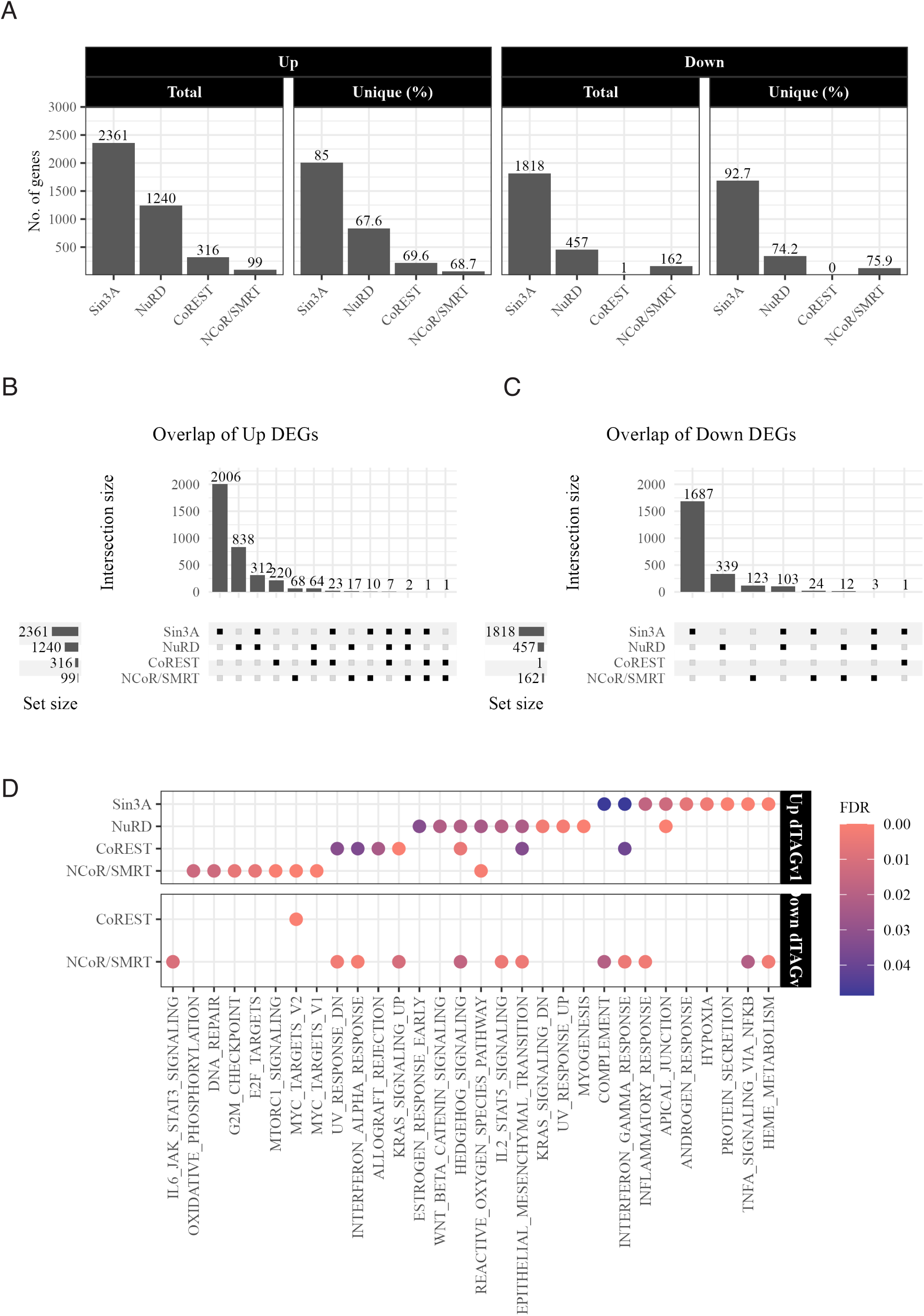
Perturbed genes are mostly complex specific with overlapping genes restricted to two complexes. A Bar plots showing the total number of significantly perturbed up and downregulated genes for each complex and the percentage of unique genes that do not overlap in the same direction with any other complex. B,C Upset plot showing intersection of up (B) and down (C) differentially expressed genes. The ‘set size’ is the total number of genes. The ‘intersection size’ is the number of genes in the indicated group. Overlaps were distinct i.e. a gene is only present in one intersection. D GSEA results for top 50 hallmark pathways. Normalised counts, filtered to only contain those with counts ≥ 50 in 3 samples, were used for input (∼12000 genes). Significant pathways (FDR <0.05) found enriched or repressed in the dTAG^V^-1 treated cells were plotted for comparison.

The NuRD complex shows the highest level of redundancy with other complexes, sharing 321 of 355, 71 of 96, and 19 of 31 upregulated DEGs, with SIN3A, CoREST and NCoR/SMRT respectively (Figure 2B). CoREST shared 31 of 96, and NCoR/SMRT 13 of 31, upregulated overlapping genes with SIN3A, whilst CoREST and NCoR/SMRT shared only 2 upregulated genes (Figure 2B). Few genes were found to overlap between three or more complexes. A similar pattern was observed for downregulated overlapping genes, with the exception NCoR/SMRT shared the majority (27 of 39) with SIN3A and not NuRD (Figure 2C).

Gene set enrichment analysis (GSEA) showed minimal pathway overlap between the complexes, consistent with the high proportion of uniquely regulated genes (Figure 2D; top). Surprisingly, given the relatively large number of downregulated genes, depletion of NuRD or SIN3A did not correlate with perturbations in any particular pathway (Figure 2D; bottom) suggesting that downregulation does not correlate with a defined physiological function. Interestingly, NCoR/SMRT depletion revealed 12 pathways in which genes are downregulated following loss of the complex. Of these, 11 pathways are upregulated following loss of other complexes (Figure 2D; bottom and Figure S3).

Taken together, these results suggest that most class 1 HDAC gene targets are complex-specific, and the complexes play largely non-redundant roles.

### Reciprocal gene regulation between complexes

The fact that depletion of NCoR/SMRT results in downregulation of genes associated with pathways upregulated by other complexes, suggests a potential reciprocal role in gene regulation. Pair-wise correlation analysis of the log_2_ fold change for all DEGs across complexes revealed a negative correlation between NCoR/SMRT with SIN3A and CoREST (-0.15 and - 0.23, respectively), and SIN3A and CoREST (-0.19) (Figure 3A). A clustered heat map of the fold change for all DEGs, further supports the finding that genes which are regulated by more than one complex are often reciprocally perturbed. In particular, SIN3A appears to regulate gene expression in an opposite manner to CoREST and NCoR/SMRT (Figure 3B).

**Figure 3.**
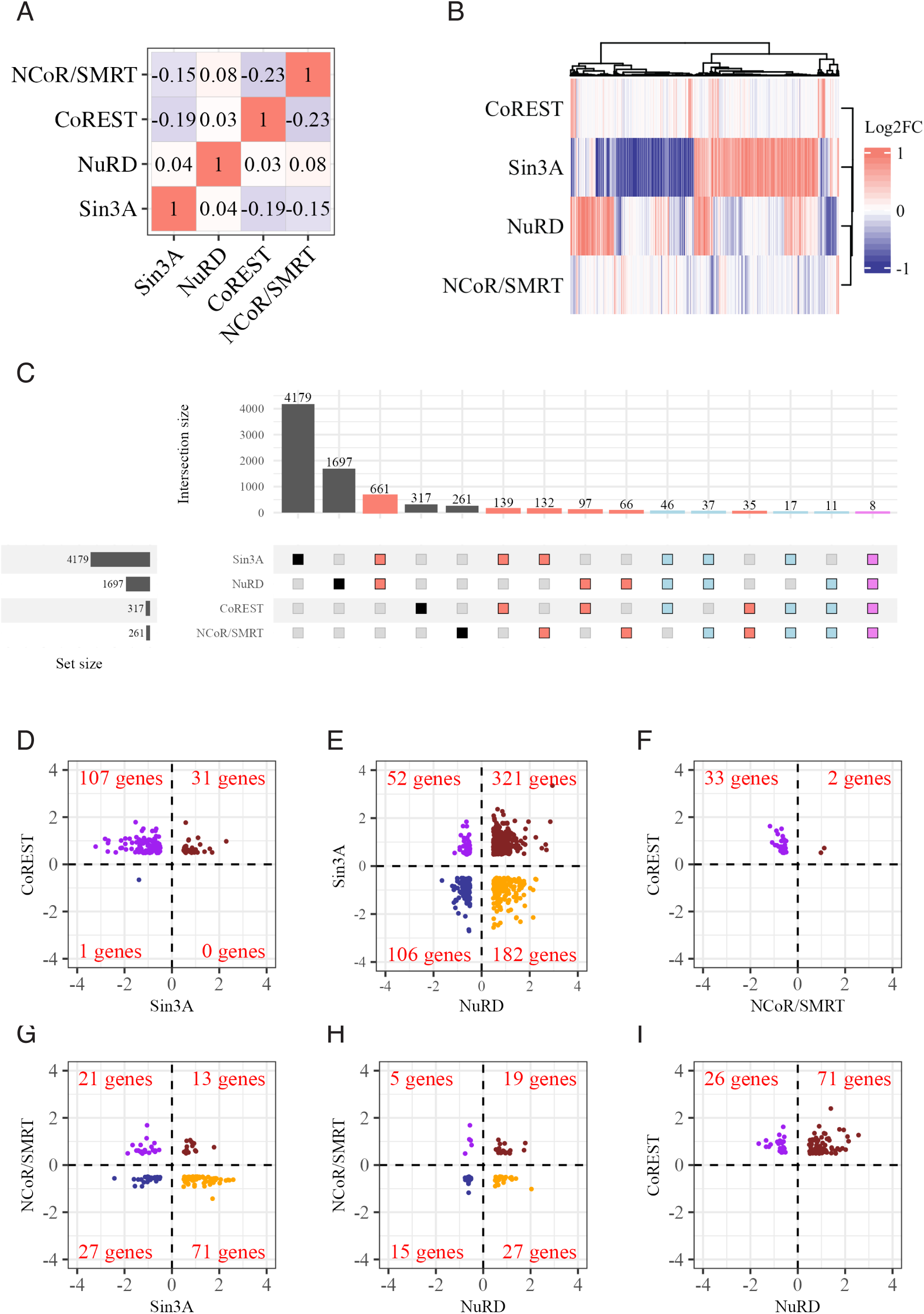
Reciprocal regulation of genes by HDAC complexes. A Correlation matrix of gene log_2_ fold change between complexes. Datasets were intersected with the total DEGs across all complexes for log_2_ fold change. Correlation was calculated using the R package stats “cor” function with Pearson calculation for correlation coefficient. B Heatmap of log_2_ fold change for all DEGs across complexes. Salmon represents positive and blue represents negative fold change. Fold change limits have been set at a maximum of 1 and -1 to aid visualization. C Upset plot showing overlap of DEGs from each complex. Coloured bars represent overlap between two (salmon), three (light blue), and four (purple) complexes. Overlaps are independent i.e. the same gene can appear in more than one overlap. This is so absolute numbers can be compared between complexes. D-I Scatter plots representing log_2_ fold change of overlapping DEGs between 2 complexes. Dark red and blue represent genes that change in the same direction whilst orange and purple represent reciprocally regulated genes.

To further explore the extent of reciprocal regulation, the overlapping DEGs from each complex were compared (Figure 3C). In each case, the total number of overlapping genes between complexes was higher than the number of genes regulated in the same direction (Figures 2B, C and Figure 3C). Notably, the SIN3A and CoREST complexes have 139 perturbed genes in common, of which 107 genes are reciprocally regulated (Figure 3C and D). Similarly, SIN3A and NuRD have 661 perturbed genes in common of which 234 show reciprocal regulation (Figure 3C and E). Reciprocal regulation was also observed between CoREST and NCoR/SMRT, and SIN3A and NCoR/SMRT (94% and 70% of genes, respectively) (Figure 3F and G). The normalised mean control counts of genes perturbed in the same or opposite directions between two complexes showed no differences based on gene expression levels (Figure S4A-F).

### NuRD and CoREST work antagonistically to Sin3A in the regulation of neuronal transcription networks

Analysis of genes perturbed by the depletion of two complexes identified few significant biological pathways. The notable exceptions are the set of genes regulated in the same direction by NuRD and CoREST; genes reciprocally regulated by SIN3A and NuRD as well as genes reciprocally regulated by SIN3A and CoREST (Figure 4A). For these gene sets, substantial overlap was observed between pathways with several involved in neuronal differentiation and development (Figure 4A).

**Figure 4.**
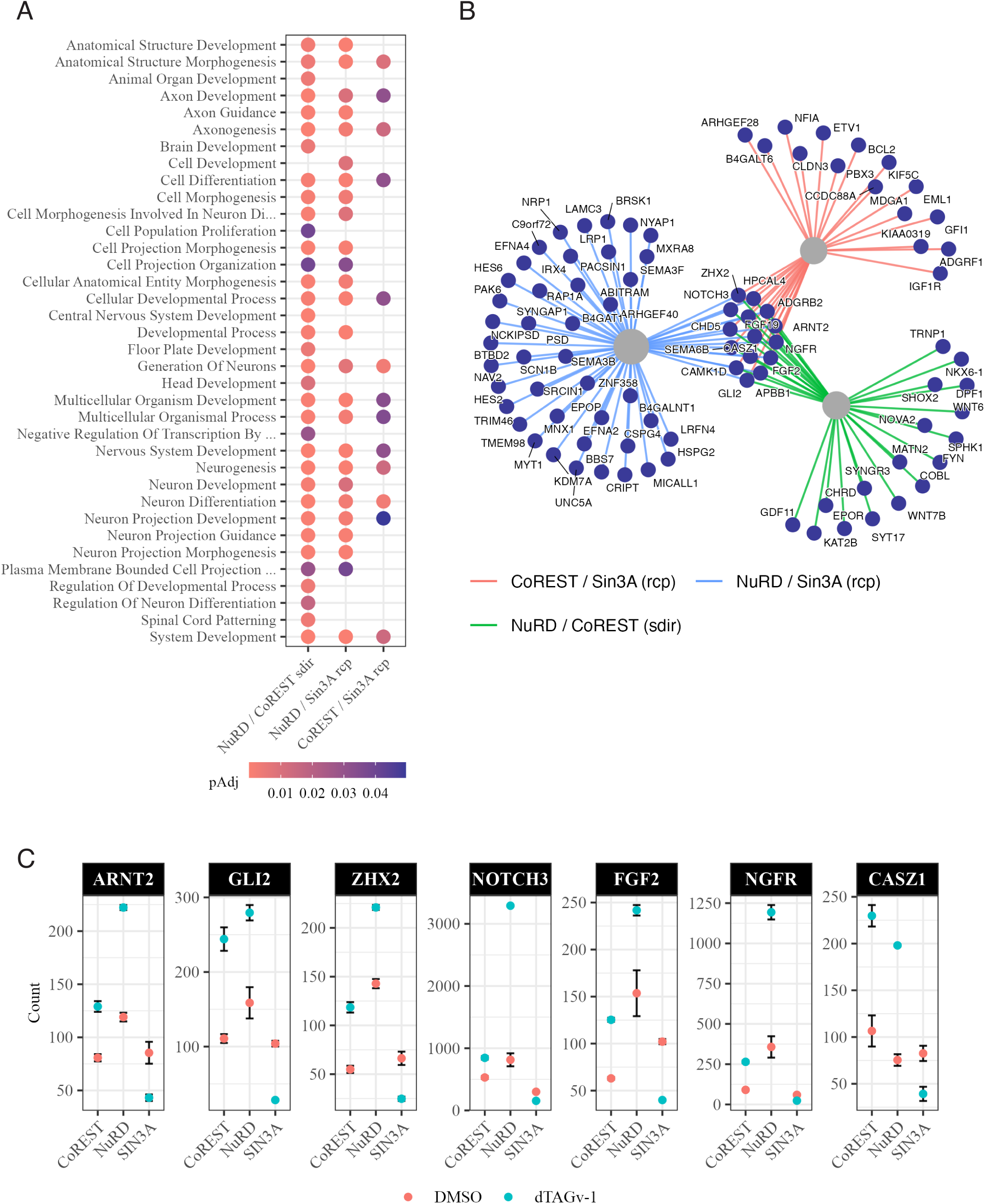
NuRD and CoREST function reciprocally to SIN3A in regulating neuronal pathways. A Significant biological process pathways enriched from overlapping genes that: change in the same direction between NuRD and CoREST, change in opposite directions between NuRD and SIN3A and change in opposite directions between CoREST and SIN3A. Pathway enrichment was performed using TopGO and adjusted p-values calculated with Benjamini & Hochberg correction using the stats package. ‘Sdir’ indicates same direction and ‘rcp’ indicates opposite direction. B Interaction map showing the significant neuronal genes. Neuronal genes from the seven overlapping neuronal pathways were used to subset the SIN3A / NuRD / CoREST overlapping gene lists. ‘Sdir’ indicates same direction and ‘rcp’ indicates opposite direction. C Normalised counts for seven genes regulated by SIN3A, NuRD and CoREST that play important roles in neuronal differentiation and/or are known targets for CoREST. Data shown are mean values with SEM, N=3.

A total of 89 DEGs were associated with neuronal pathways across the three gene sets, with 32 changed in the same directions by CoREST and NuRD, 57 reciprocally regulated by NuRD and SIN3A, and 30 reciprocally regulated by CoREST and SIN3A (Figure 4B). In general, genes were downregulated following SIN3A, and upregulated following NuRD or CoREST, depletion (Figure S5). Of the 15 shared genes, seven - NOTCH3, ZHX2, GLI2, ARNT2, FGF2, NGFR and CASZ1 - are key proteins involved in neuronal development and differentiation (Becquart *et al*, 2020; Bruno *et al*, 2023; Liu *et al*, 2023; Radecki *et al*, 2020; Rusanescu & Mao, 2014; Woodbury & Ikezu, 2014; Wu *et al*, 2009) and are known CoREST targets (Egolf et al 2019). Except for NOTCH3, the normalised control counts for these genes are all less than 200 counts and increase following CoREST and NuRD depletion and decrease following SIN3A depletion (Figure 4C).

This antagonistic relationship suggests CoREST and NuRD work cooperatively to repress gene expression associated with neuronal processes, counteracting activation by SIN3A. In response to external signals, this repression maybe lost, allowing SIN3A to activate gene expression and drive differentiation forward.

### Transcription factor regulation is not associated with gene expression changes at 6 h

We used HOMER analysis to identify transcription factor binding sites (TFs) associated with particular DEGs. This showed enrichment for specific TFs in the SIN3A, NuRD and CoREST significant gene lists with minimal overlap between complexes (Figure 5A). Genes upregulated following CoREST depletion were associated with SNAIL1 and SLUG - hallmark mesenchymal TFs known to recruit CoREST and regulate gene expression (Ferrari-Amorotti *et al*, 2013; Lin *et al*., 2010a; Lin *et al*, 2010b; Wu *et al*, 2012). Seven of the same TFs were found enriched between SIN3A and NuRD DEGs, although four of these were associated with gene expression changes in opposite directions (Figure 5A).

**Figure 5.**
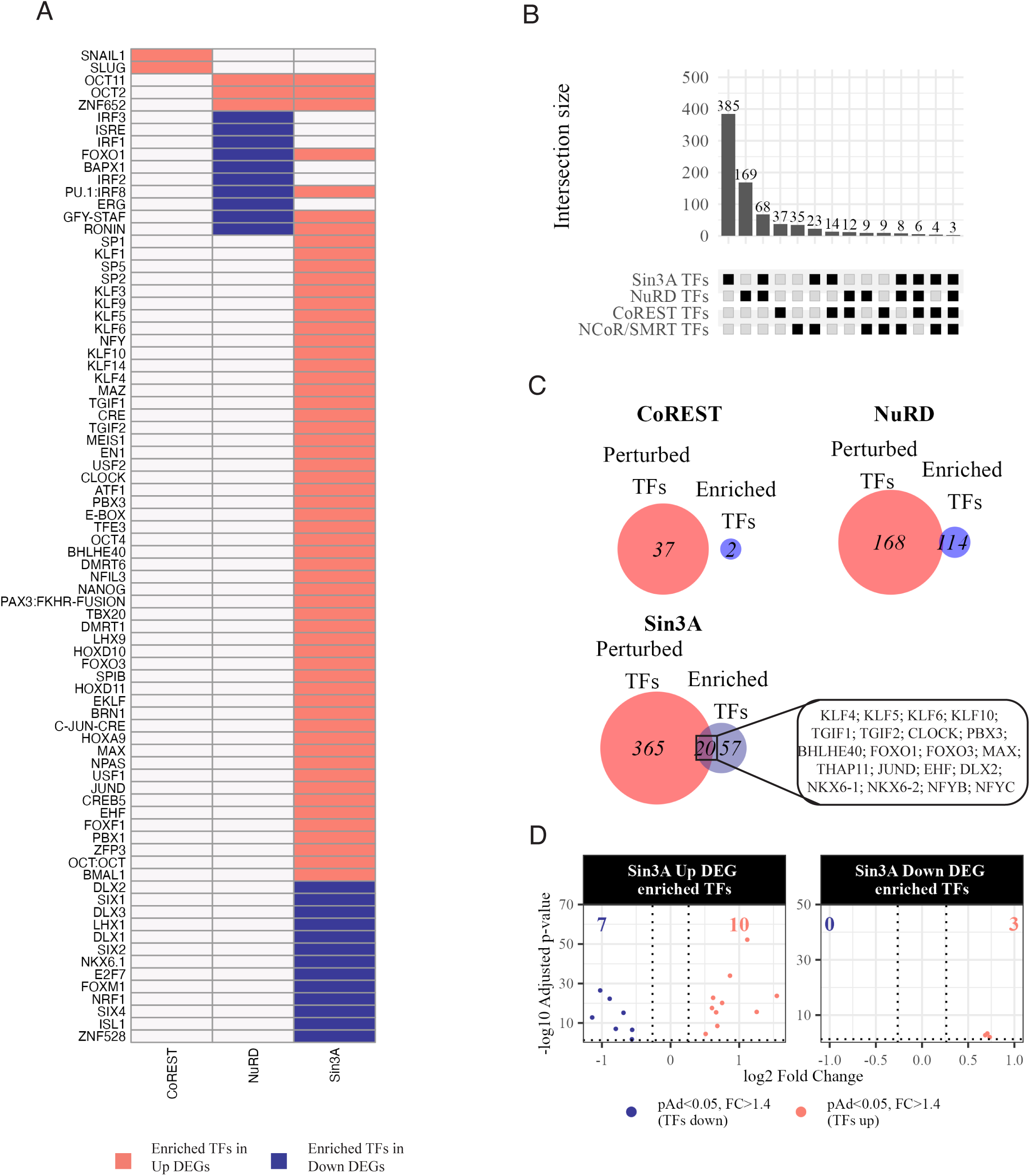
Perturbation of transcription factor expression and identification of enriched transcription factor gene targets. A HOMER results for transcription factors with enriched binding sites for significant up and down DEGs for CoREST, NuRD and Sin3A. B Intersection of identified perturbed transcription factors from each complex dataset. Overlaps shown are independent of each other, i.e. the same gene can appear in more than one overlap. This is so absolute numbers can be compared between complexes. C Overlap of differentially expressed transcription factors and those associated with perturbed genes for CoREST, NuRD and SIN3A. The overlapping transcription factors for SIN3A are also identified. D Log_2_ FC vs adjusted p-value of transcription factors both perturbed and associated with either up (left) or down (right) SIN3A DEGs.

To determine if perturbations in gene expression of the enriched TFs could account for changes in expression of gene targets, DEGs were overlapped with known human transcription factors (Figure S6A and B) (Lambert et al 2018). Overlapping TFs between complexes followed the same pattern seen with the total DEGs, with only eight regulated by three complexes and just three (ARNT2, NKX6-1 and ZHX2) regulated by all four (Figure 5B and S6C). Pairwise comparisons revealed similar patterns to total DEGs overlaps (Figure S7A-F), with a notable reciprocal relationship between TFs regulated by CoREST and NCoR/SMRT.

Overlap of TFs perturbed by CoREST and NuRD with those identified from HOMER analysis revealed little overlap (0 and 1, respectively), suggesting changes in TF gene expression does not account for perturbations in gene targets but is rather recruitment to TFs (Figure 5c). Following SIN3A depletion, 20 TFs associated with perturbed genes also showed changes in gene expression; with several of the same family members affected (KLF4/5/6/10; TGIF1/2; FOXO1/3; NKX6-1/6-2) (Figure 5C). The majority of these (17 of 20) were associated with upregulated genes with SIN3A depletion having both activator and repressive effects on TF gene expression (Figure 5D). Both KLF and TGIF families are known to be SIN3A interactors (Kwon *et al*, 2016; McConnell & Yang, 2010) that could suggest changes in their expression represent a compensatory mechanism for SIN3A loss, and changes to gene targets are still a direct consequence of SIN3A depletion.

## Discussion

This is the first study to use a PROTAC approach to induce rapid, complex-specific depletion of six endogenous class 1 HDAC complexes – SIN3A, NuRD, CoREST, MiDAC, MIER and NCoR/SMRT. Overall, tagging endogenous genes within each complex had minimal effect on protein expression and no effect on complex enzyme activity. Knockout of MTA1 and MTA3, along with tagging MTA2, from the NuRD complex, did result in somewhat reduced cell-proliferation. However, gene-editing the other complexes had no discernible proliferation defect. Treatment with the dTAGv1 PROTAC resulted in >95% degradation of the target protein within 1 h.

Changes in the transcriptome at 6 h following depletion of HDAC complex proteins are the earliest measured to date. This has allowed identification of primary targets for six class 1 HDAC complexes and enabled comparative analysis of gene targets between complexes using the same model system. Depletion of SIN3A, NuRD, CoREST or NCOR/SMRT perturbed the expression of 6,461 genes, more than 50% of all genes ‘expressed’ in HCT116 cells. Even this may be an underestimate of class 1 HDAC complex gene targets, since we did not target the SIN3B or RERE complexes which are thought to have distinct functions (Adams *et al*., 2018; Kumar & Wang, 2016; Li *et al*, 2024).

Over two-thirds of altered genes were only perturbed when a single complex was depleted, suggesting a predominantly non-redundant role in cells. Only 508 genes were regulated by more than one complex in the same direction and largely restricted to two individual complexes with NuRD sharing the most overlap with SIN3A, CoREST and NCoR/SMRT.

Whether these overlaps are of a redundant or cooperative nature is difficult to distinguish. On the one hand, it is possible two complexes physically interact to regulate gene expression, as is reported for LSD1 and NuRD (Wang *et al*, 2009b), LSD1 and SIN3A (Yang *et al*, 2018), and LSD1 and DNTTIP1 (Liu *et al*., 2022). Both are required to alter gene expression and depletion of either has a similar effect on gene expression. Alternatively, two complexes could function independently on the same gene, targeting different epigenetic marks. In this case, knockdown of both may result in a cumulative change in gene expression. Further experiments targeting two complexes in the same cell line that can be degraded either independently or together would be an interesting approach to further understand these relationships.

What is most striking is the finding that of 1130 genes regulated by any two complexes, 524 were of a reciprocal nature. Notably, CoREST and NuRD work cooperatively, or redundantly, to repress genes associated with neuronal development and differentiation while competing with SIN3A which is activating the same genes. This is in apparent contrast with several studies showing SIN3A and CoREST maintain neuronal gene repression in non-neuronal cells through interaction with REST (Grimes *et al*, 2000; Jayaprakash *et al*, 2021). One explanation is that the genes identified in this study are involved in non-neuronal pathways that are activated in response to external signals. For example, several genes identified - NOTCH3, FGF2, NGFR and CASZ1 – have been shown to be upregulated following LSD1 inhibition and can drive epithelial differentiation (Egolf *et al*, 2019).

The observation of several reciprocal relationships between complexes provides further evidence supporting the importance of developing complex specific inhibitors. Current FDA approved HDAC inhibitors have limited use in the clinic due to severe side effects or limited efficacy due to targeting multiple HDAC isoforms/complexes (Bondarev *et al*, 2021; Huang *et al*, 2024; Millard *et al*., 2017). These results may provide an explanation as to why this is the case: not only will inhibition of all complexes perturb multiple non-redundant pathways, but also if disease pathways are regulated antagonistically by two complexes, the effect of an inhibitor may be reduced.

Taken together, these results are the first to provide a comparative analysis of primary gene expression changes for six class 1 HDAC complexes. It is particularly striking that just 4 complexes alter the expression of >6000 genes. Yet this is not simply because HDAC complexes have a non-specific role in suppression of gene activity since there is very little redundancy between complexes suggesting a remarkable specificity of gene regulation. Although some overlap was observed, this was largely restricted to between two complexes and ∼50% of all shared genes showed a reciprocal relationship. This highlights the need for further understanding of the gene regulatory roles of each complex in a cell type specific manner and the development of complex specific inhibitors to improve efficacy and safety.

## Methods

### Cell culture and proliferation

HCT116 colon cancer cells were cultured in Dulbecco’s modified eagle’s media (DMEM) supplemented with 10% foetal bovine serum (FBS) and 1% penicillin, streptomycin, glutamate (PSG). Cells were detached from plates using TryplE (Invitrogen®). CRISPR/Cas9 engineered HCT116 lines were cultured in the same way as the parental line. Cells were passaged for a maximum of four weeks to minimise further genetic divergence between modified cell lines. Cells were treated with a final concentration of 100 nM dTAG^V^-1 (R&D Systems) for indicated time points. An equal volume of DMSO was added to cells as a vehicle control. To measure cell growth, 0.15x10^6^ cells were seeded into 6-well plates and counted every 24 h using a BoiRad automated cell counter.

### CRISPR/Cas9 targeting of endogenous alleles

HCT116 colon cancer cells were selected due to their stable haplotype, previous successful CRISPR/Cas9 targeting and terminally differentiated state. Initial RNA sequencing on the parental HCT116 cells confirmed expression of all HDAC complex components and no effect of the protacs dTAG-13 or dTAG^V^-1 on gene expression (Figure S2A and B). Normalised counts for dTAG-13 and dTAG^V^-1 associated E3 ligases suggested VHL is expressed ∼4-fold higher than CRBN and so dTAG^V^-1 was used for all experiments (Figure S2C).

The unique complex components SIN3A (Sin3), MTA1/2/3 (NuRD), KDM1A (CoREST), DNTTIP1 (MiDAC), MIER1/2/3 (MIER) and HDAC3 (NCoR/SMRT) were selected for CRISPR/Cas9 targeting based on gene expression and presence of paralogues (Figure S2A). SIN3B, MTA1 and RERE was also targeted for knock in but were unsuccessful. All genes were targeted at the C-terminus apart from MIER1, which was N-terminally targeted (Ran et al 2013; Natsme et al 2016). Guide RNAs (gRNAs) and homology arms were selected using ‘CRISPR design and analyse guides’ tool on Benchling (https://www.benchling.com/). A list of oligonucleotides for guide RNA (gRNA) duplexes and PCR amplification of donor templates are provided in Table S1.

CRISPR/Cas9 gene blocks (gblocks) for knock in (KI), with DNA sequences encoding FKBP12^F36V^ and 3x FLAG flanked by homology arms (donor template), and gRNAs for both KI and knock out (KO) targeting were synthesized by IDT. The length of homology arms varied between 350bp and 500bp. For MTA2, KDM1A, DNTTIP1 and MIER1 KI cell lines, gblocks were PCR amplified and cloned using TOPO TA Cloning Kit (Thermo Fisher). Duplex gRNAs were cloned into the BbsI site of PX330-U6-Chimeric_BB-CBh-hSpCas9. The donor template and gRNA/Cas9 plasmids were co-transfected using jetPRIME transfection reagent (Polyplus). Except for MIER1, antibiotic resistant markers were inserted in-frame and transfectants selected using double antibiotic selection (Geneticin and Hygromycin B). For MIER1, single cells expressing Cas9-mCherry were FACS sorted.

CRISPR/Cas9 KO of MIER2/3 and MTA1/3 genes was performed in MIER1 and MTA2 tagged cell lines like KI except gRNA duplexes were cloned into BbsI site of pSpCas9(BB)-2A-Puro V2.0 vector, transfected and selected using Puromycin for 48 h.

CRISPR KI gene targeting for HDAC3 and SIN3A was performed as previously described (Nabet 2018). Briefly, 30-nucleotide microhomology arms were inserted in primers to amplify two cassette sequences encoding tags and Puromycin and Blasticidin genes. The rest of the protocol was as described above.

Following antibiotic selection or FACS, single colonies for KI or KO were expanded in antibiotic-free media and targeting confirmed using Sanger sequencing. Successful targeting was confirmed at the protein level by western blotting (Figure S8A). Due to poor antibodies, deletion of MIER2 protein could not be shown, but sequencing confirmed deletion in the targeted exon, like MIER3, that would result in a frameshift (Figure S8B).

### Isolation of cytoplasmic and nuclear proteins

Cytoplasmic and soluble nuclear proteins were isolated from cell pellets using Abcam kit (ab219177; now discontinued) following the manufacturer’s instructions. Briefly, cell pellets were resuspended to homogeneity in cytoplasmic extraction buffer supplemented with protease inhibitor cocktail (PIC). Samples were vortexed at max speed for 10 s before incubation on ice for 10 min and a further 10 s vortex. Nuclei were pelleted by centrifugation at 1000 rcf for 3 min at 4 °C. The nuclear pellet was resuspended in soluble nuclear buffer supplemented with PIC, vortexed for 10 s and incubated on ice for 15 min with a 10 s vortex every 5 min. Precipitated DNA and insoluble material was removed by centrifugation at 5000 rcf for 3 min at 4 °C. The supernatant containing soluble nuclear proteins was collected for downstream applications. Protein concentration was determined using the BioRad protein assay dye following manufacturer’s instructions.

### Western blotting

Where applicable, sample concentrations were normalised before separation using NuPAGE 4-12% Bis-Tris gels (Invitrogen®) in 1x MES SDS running buffer. Proteins were transferred to nitrocellulose membranes using a semi-dry transfer system with everything pre-soaked in transfer buffer (48 mM Tris, 39 mM glycine, 20% methanol, pH 9.2) at 75 mA /per gel for 1.5 h. Membranes were air-dried at room temperature for 30 min before blocking for 1 h in tris-buffered saline (TBS; 20 mM Tris, 150 mM NaCl, pH 7.6) with 0.5% Tween-20 (TBS/T) and 5% milk. Membranes were incubated with primary antibody (Table S2) overnight at 4 °C and secondary antibodies (IRdye 680RD goat anti-mouse or IRdye 800CW goat anti-rabbit (both 1:20000); Licor) for 1 h at room temperature. All antibodies were diluted in blocking buffer. Membranes were image using an Odyssey Clx imager and processed using image studio (v5.5.4).

### Immunoprecipitation

Complexes were immunoprecipitated from the soluble nuclear fraction. Briefly, pre-washed anti-mouse dynabeads^®^ or protein A Dynabeads^®^ (1.2 mg; Invitrogen®) were incubated overnight at 4 °C with antibodies (mouse monoclonal M2 anti-FLAG (10 μg; Merck) and rabbit anti-LSD1 (2 μg; Abcam)) to bind. Antibody bead complexes were washed before addition to the soluble protein isolate and incubated overnight at 4 °C with end over end rotation. The following day, the beads were washed x1 in soluble nuclear buffer and x2 in assay specific buffer for downstream applications. All washes were carried out using a magnet (Invitrogen®).

### HDAC assay

HDAC activity was measured as described previously (Watson et al 2012) using the HDAC Assay Kit (Active Motif). FLAG-labelled endogenous protein complexes were immuno-precipitated from ∼50x10^6^ cells using the M2 FLAG antibody. The FLAG immuno-precipitate from the parental HCT116 line was used as a control in this assay. Complexes were kept on beads and incubated with 100 μM substrate for 1 h at 37 °C before addition of an equal volume developing/stop solution. Fluorescence was measured using a Victor X5 plate reader (PerkinElmer).

### Demethylase assay

Following immunoprecipitation with anti-FLAG or anti-LSD1 antibodies, protein-bead complexes were washed x2 with assay buffer (20 mM HEPES/KCl pH 7.5, 50 mM NaCl, 0.1 mg/ml BSA), resuspended in assay buffer and added, in duplicate, to 96 well black plates (Corning). A solution of 0.4 mg/ml HRP and 100 μM Amplex™ Ultrared (final concentrations: 0.04 mg/ml HRP, 10 μM Amplex™ Ultrared) was added followed by the H3K4Me peptide (final concentration (15 μM) to the side of the well. The plate was covered, briefly centrifuged and fluorescence measured in a Victor X5 plate reader (PerkinElmer) over 21 min with an excitation wavelength of 530 nm and emission wavelength of 590 nm. For this study, a FLAG-IP from the parental HCT116 cell line was used as a control.

### RNA sequencing

To identify primary changes in gene expression elicited upon complex knockdown, RNA sequencing was performed after 6 h dTAG^v^-1 treatment. We chose 6 h as this seemed an appropriate length of time to allow for changes to occur (e.g. target degradation, histone modification, chromatin remodelling, sufficient change in mRNA level).

Cells were seeded into 6-well plates and allowed to settle for 24 h before treatment with 100 nM dTAG^V^-1 or DMSO for 6 h. Cells were washed with PBS, detached using TrypLE, diluted in PBS and transferred to nuclease-free 1.5 ml Eppendorf’s. Cells were collected by centrifugation at 400 rcf for 5 min at room temperature, the supernatant decanted and total RNA isolated by robot and quality checked using the Genomics facility at the University of Leicester. Each experiment was carried out in triplicate.

RNA sequencing was performed by Novogene using a NovaSeq 6000 S4 or NovaSeq X plus platform using 150 base-pair pair-end sequencing to a depth of 20 million reads. The resulting sequencing files were quality checked using fastqc ((Andrews, 2010), v0.12.1). Sequencing files were aligned using hisat2 ((Kim *et al*, 2019), v12.3.0) with standard options to the human genome build GRCh38 release 111 (Dyer *et al*, 2025). Resulting files were converted to binary alignment map (BAM) format, sorted by genomic coordinate and indexed using samtools ((Li *et al*, 2009), v1.17) with standard options. Count files were made using LiBiNorm ((Dyer *et al*, 2019), v2.5) (options: -z -r pos -I gene_name -s reverse) and the GRCh38 release 111 gene transfer format file.

Unless stated otherwise, analysis was performed using R (v4.4.1). DESeq2 (Love et al 2014; v1.44.0) was used to perform differential expression (Wald test with local fit type and the Benjamini-Hochberg method for P-value correction). The package apeglm (Zhu et al 2018; v1.26.1) was used to provide shrinkage estimates for log fold changes. Genes with counts less than 50 in 3 or more samples were excluded. Gene set enrichment analysis (GSEA) (Subramanian et al 2005; Mootha et al 2003; Liberzon et al 2015) was tested against the msigdb hallmark gene set. Homer (Heinz et al 2010; v5.0.1; options: gene set permutations, FDR < 5%) was used to test gene sets for transcription factor enrichment. Enrichment for biological processes was performed using the R package TopGO ((Alexa, 2024), v2.56.0) with correction for multiple comparisons calculated using the stats package (v4.4.1) using the Benjamini & Hochberg method. The R package marge ((Amezquita, 2024), v0.0.4.999) was used to import homer results into R and the package HGNChelper ((Oh *et al*, 2020), v0.8.1) used to check HGNC symbols and correct to current version.

## Additional R programs

Data was visualised using the following R programs: ComplexHeatmap ((Gu *et al*, 2016), v2.20.0), ComplexUpset ((Lex *et al*, 2014), v1.3.3), enrichplot ((Guangchuang, 2024), v1.24.4), ggcorrplot ((Kassambara, 2023), 0.1.4.1), ggh4x ((van den Brand, 2024), v0.2.8), ggplot2 ((Wickham, 2009), v3.5.1), VennDiagram ((Chen & Boutros, 2011), v1.7.3).

## Data availability

The RNA sequencing datasets produced in this study are available at Gene Expression Omnibus and will provided upon publication.

## Acknowledgements

We are grateful to Dr. Nicolas Sylvius and the NUCLEUS genomics facility for help with RNA extractions and assessing RNA quality. This research used the ALICE High Performance Computing facility at the University of Leicester.

