## Supplementary Information for "Rapid degradation of 6 class I HDAC complexes reveals minimal functional overlap between complexes"

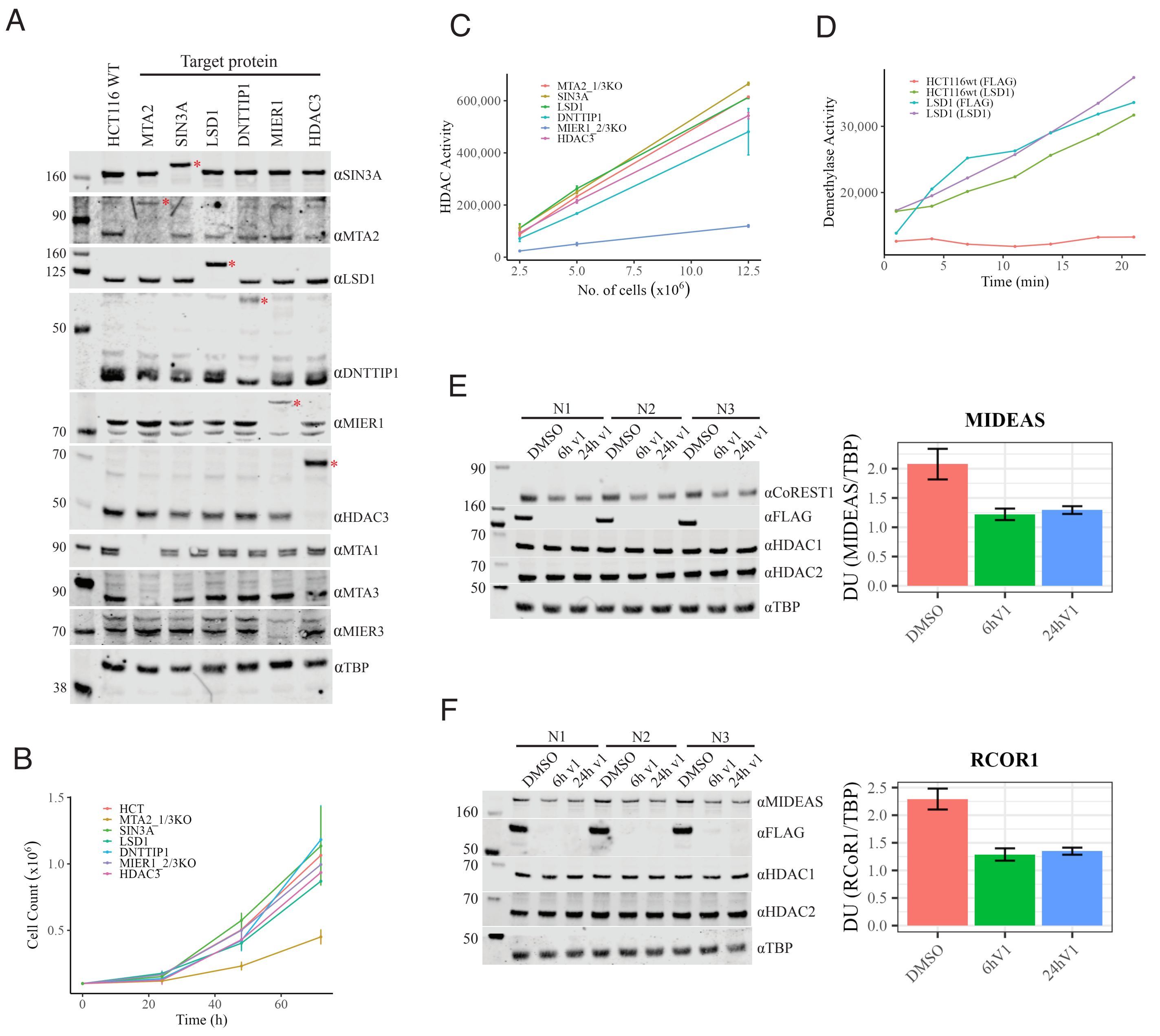


**Figure S1. CRISPR targeting endogenous proteins does not affect protein expression proliferation or enzymatic activity.**

A Western blot showing expression of CRISPR targeted proteins in relation to HCT116 cell lines with other targeted proteins and parental HCT116 cells. The CRISPR targeted protein is identified (*). TATA-binding protein was used as a loading control.

B Cell proliferation over 72 h of each targeted HCT116 cell line and parental HCT116 cell line. Results are mean±SEM (n=3).

C HDAC activity of FLAG-immunoprecipitated complexes. Technical duplicates were carried out using the immunoprecipitate from the indicated number of cells. The HDAC activity of FLAG-immunoprecipitated parental HCT116 cells was subtracted from values as background.

D Demethylase activity of targeted LSD1 protein. FLAG- and LSD1-immunoprecipitated complexes from parental HCT116 cells and targeted LSD1 HCT116 cells were used to measure histone peptide demethylation over 21 minutes.

E Western blot of targeted LSD1 HCT116 cells following incubation with 100 nM dTAG$^{V}$-1 for 6 h or 24 h. Quantitaiton of signal for RCoR1 is shown in the graph (n=3).

F Western blot of targeted DNTTIP1HCT116 cells following incubation with 100 nM dTAG$^{V}$-1 for 6 h or 24 h. Quantitaiton of signal for MIDEAS is shown in the graph (n=3)

Data informaiton: In (B,C,E,F), data are presented as mean ± SEM.

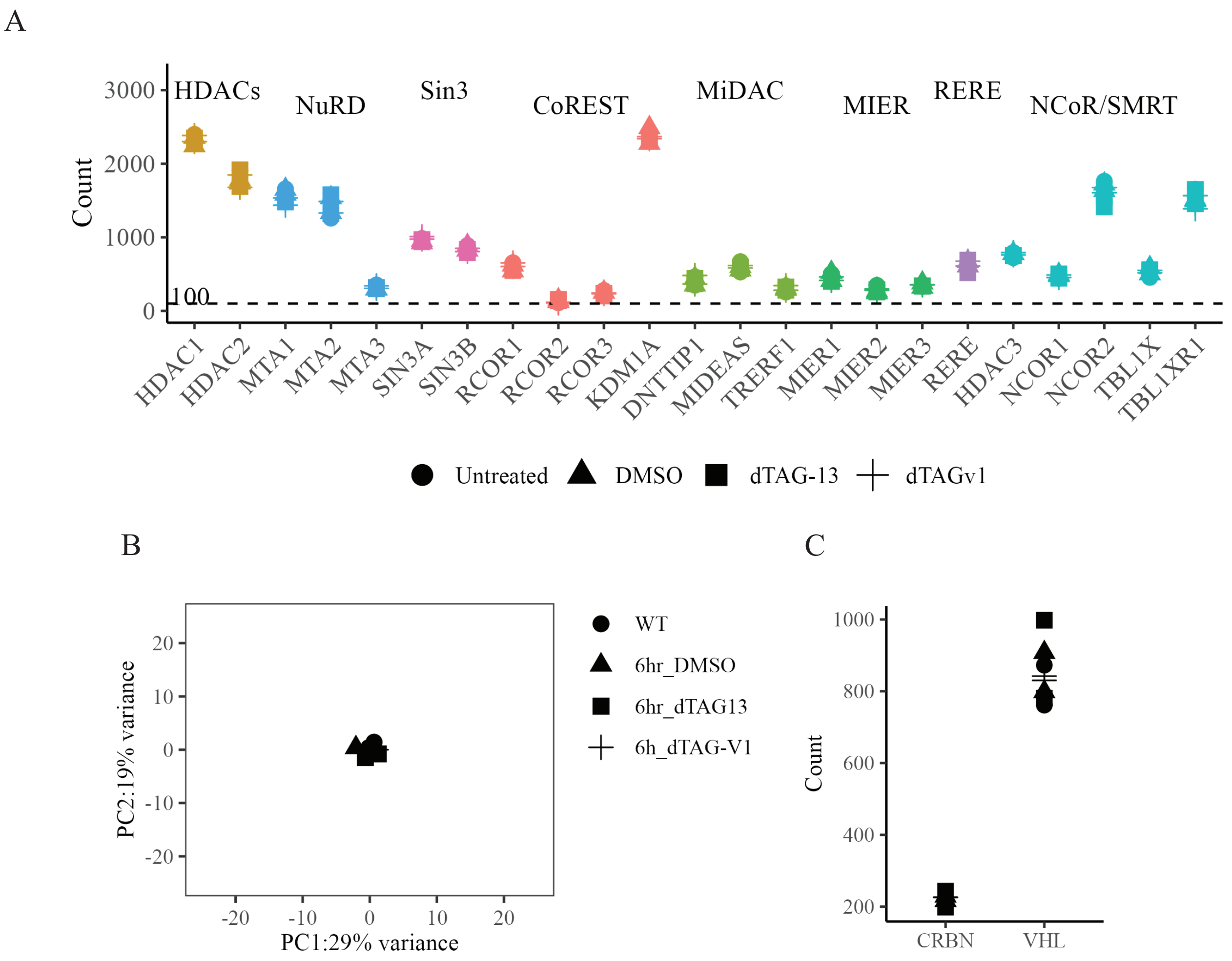


**Figure S2. DTAG-13 and dTAG$^{V}$-1 have no effect on gene expression of HDAC complex proteins**.

A The HCT116 cell line was treated for 6h with 100 nM dTAG-13 or dTAG$^{V}$-1 before harvesting and RNA-sequencing. Normalised counts for untreated, DMSO (control), dTAG-13 and dTAG$^{V}$-1are shown.

B Principal Componant Analysis of RNA-sequencing results from HCT116 cells following control or PROTAC treatments.

C Normalised counts for dTAG-13 target E3 ligase CRBN and dTAGV-1 E3 ligase VHL in HCT116 cells

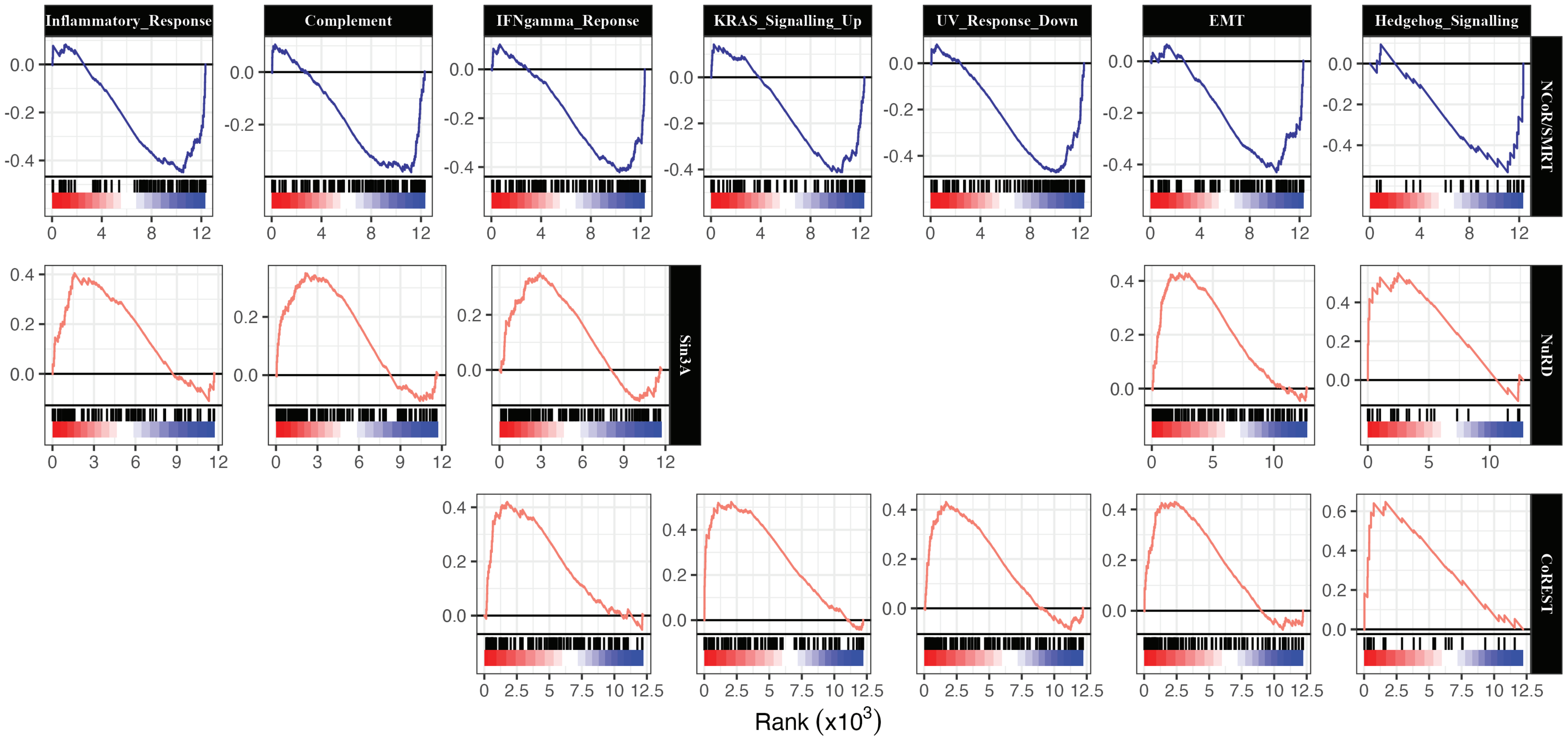


**Figure S3. NCoR/SMRT shows reciprocal association with several GSEA pathways identified in other complexes.**
GSEA graphs representing the seven pathways repressed after NCoR/SMRT depletion that are enriched following either SIN3A, NuRD or CoREST depletion. EMT – Epithelial Mesenchymal Transition.

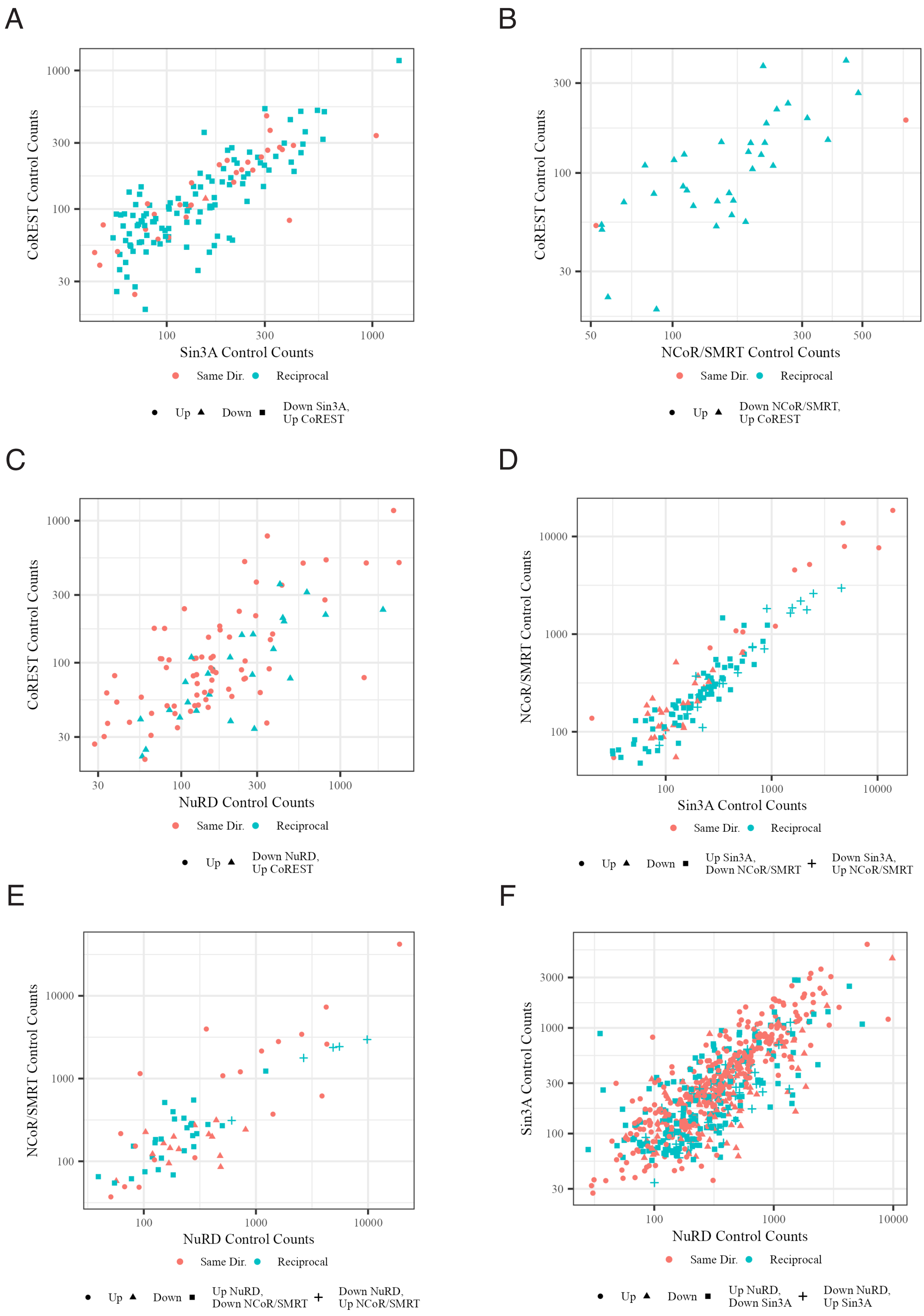


**Figure S4. Gene changes in the same or reciprocal direction between two complexes do not show differences in expression.**

A-F Normalised mean control counts for overlapping genes that change in either thesame or opposite directions between two HDAC complexes. Symbol colour donates the same (salmon) and opposite (light blue) direction while symbol shape indicates whether the genes change up or down for each complex. The specific complexes are stated on the x- and y-axis.

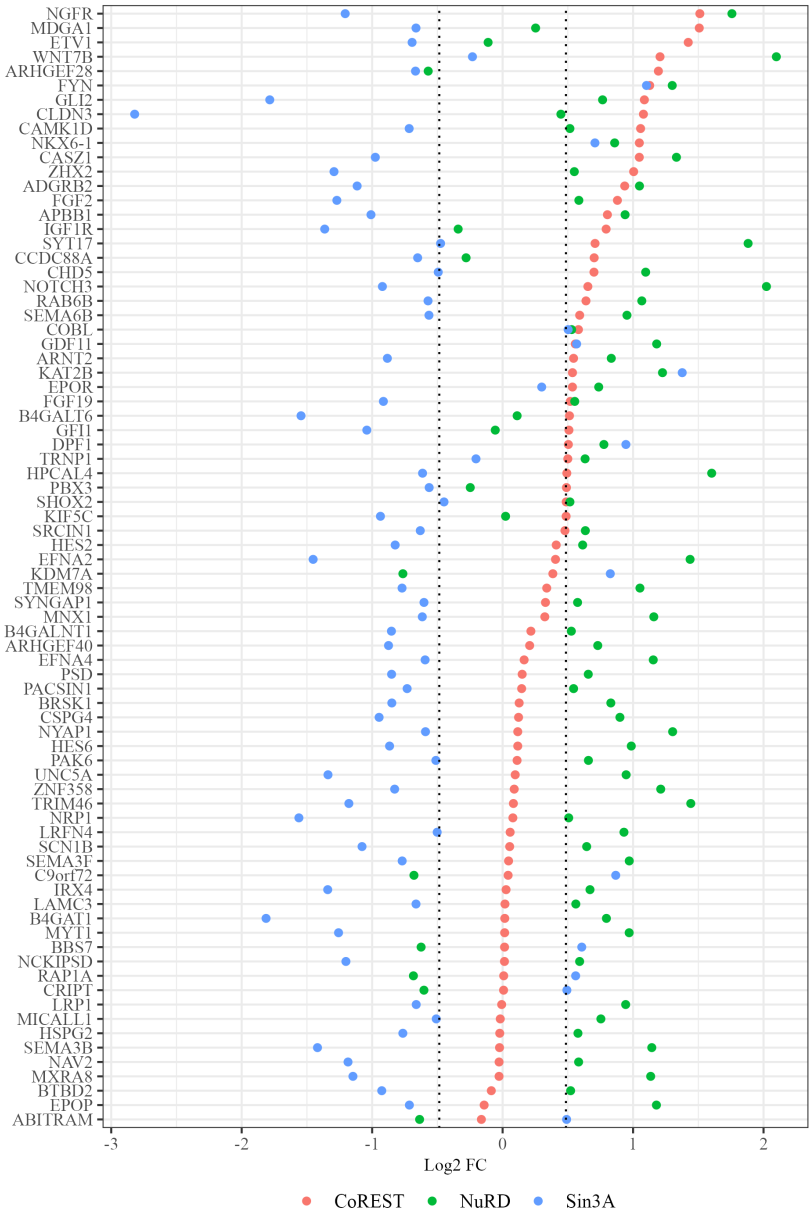


**Figure S5. CoREST and NuRD repress while SIN3A activates neuronal gene expression in HCT116 cells.**
Log2 FC of the 78 genes associated with overlapping neuronal pathways present in SIN3A, NuRD and CoREST datasets. Genes are ordered based on calculated log2 fold change in the CoREST dataset. Dotted lines at ± 0.485 represents the 1.4-fold change used to select significant genes in this study.

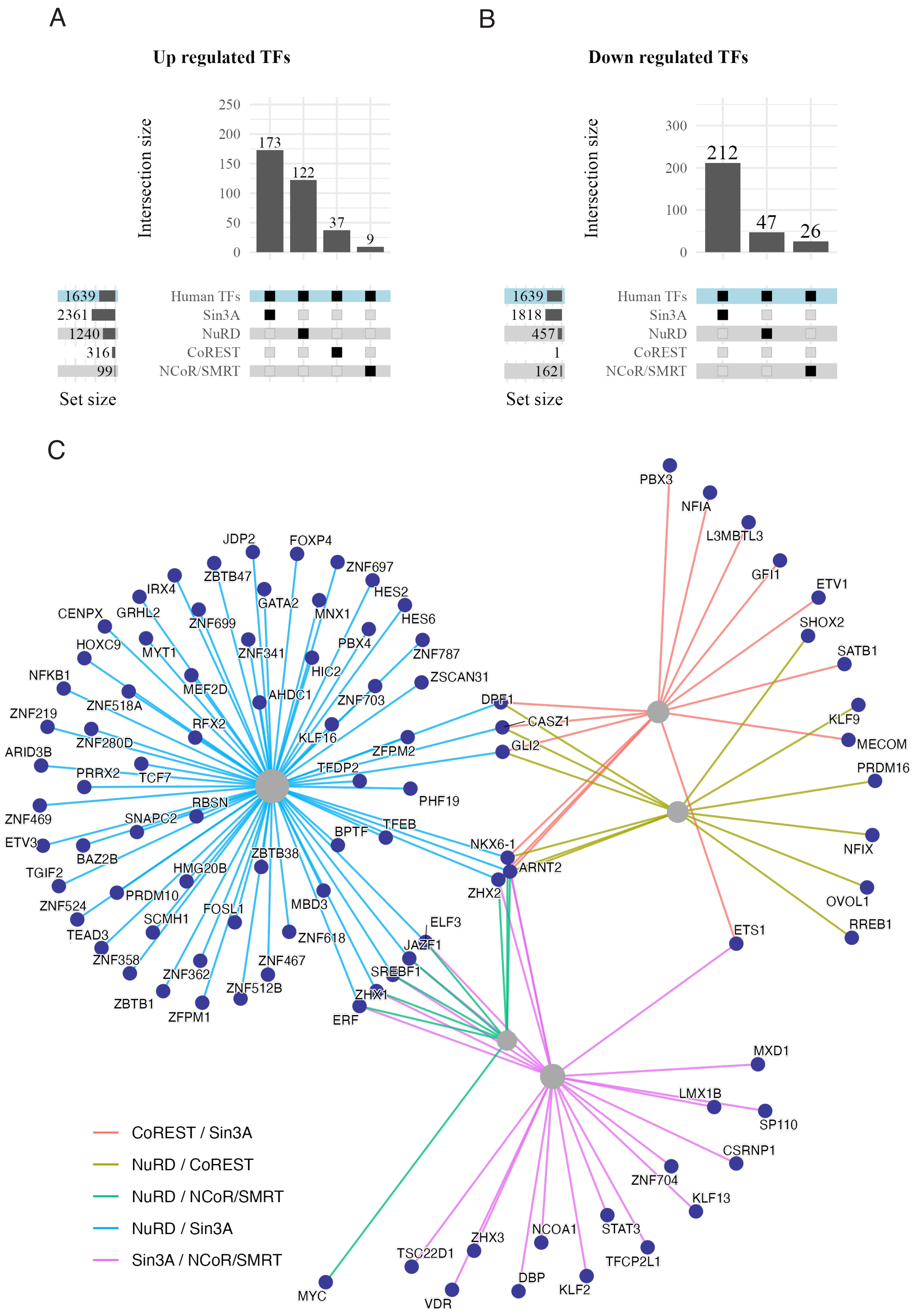


**Figure S6. Perturbed transcription factors that overlap are restricted to between two HDAC complexes**

A,B Number of significant up (A) and down (B) DEGs from each complex that are known human transcription factors. Overlap was performed using a human transcription factor list (Lambert et al 2018).

C Interaction map showing transcription factors that overlap between two complexes and identifying those that are perturbed by more than two complexes.

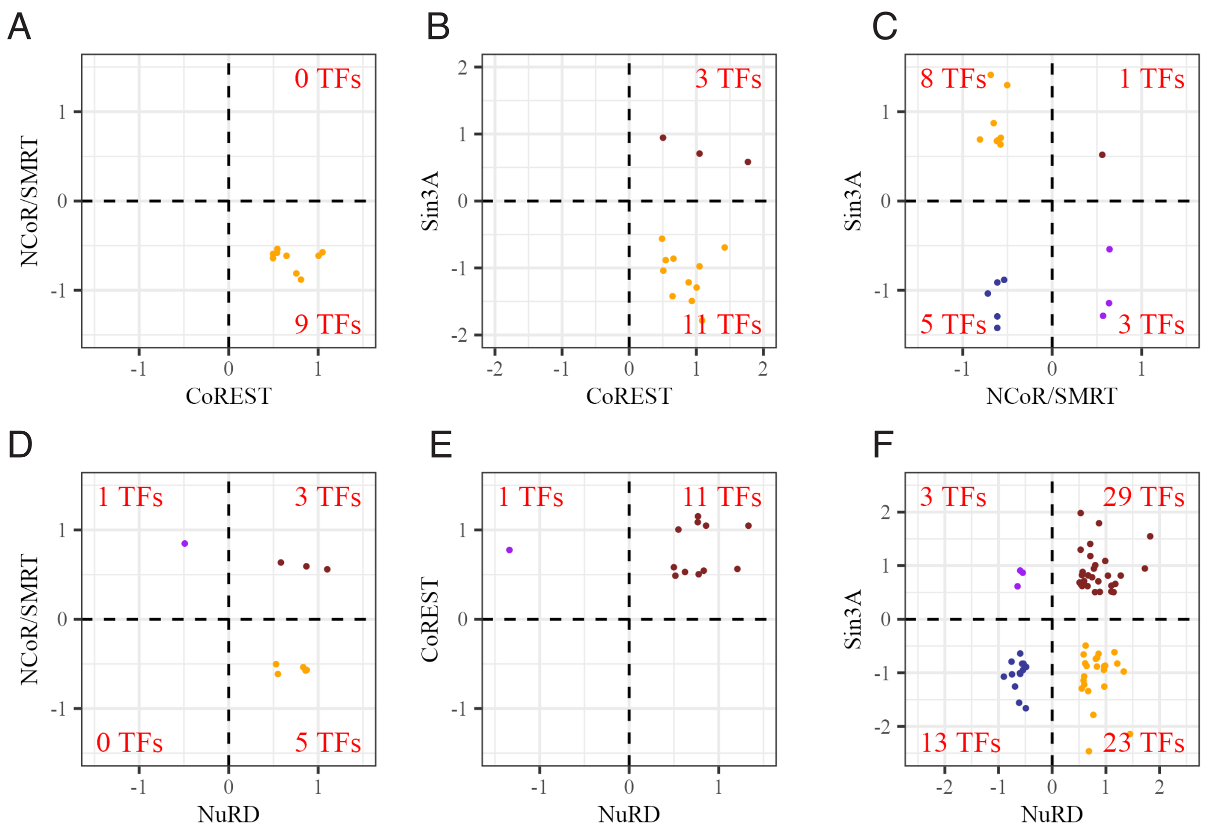


**Figure S7. Reciprocal regulation of transcription facter gene expression between complex pairs.**

A-F Scatter plots representing Log2 fold change of overlapping perturbed transcription factors (TFs) between pairs of complexes. Dark red and blue represent TFs that change in the same direction whilst orange and purple represent reciprocally regulated TFs.

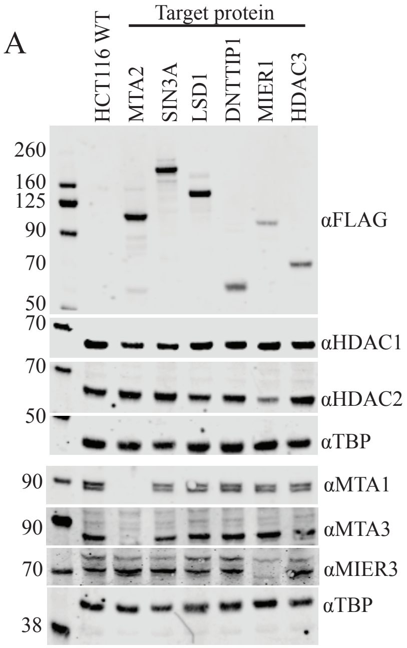


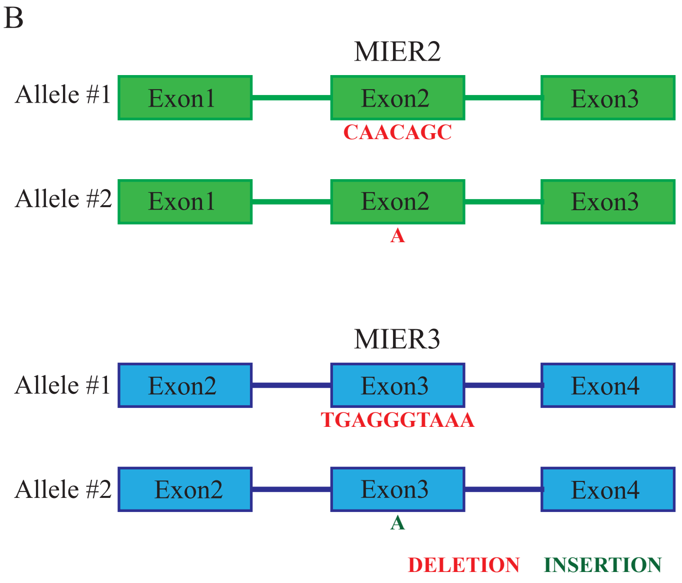


**Figure S8. Conformaiton of successful CRISPR targeting**

A Western blot showing of CRISPR-targeted proteins in HCT116 cells and parental HCT116 cells.

B Graphical representation of sequencing results following targeting of MIER2 and MIER3 for knockout.

**Table S1. Sequences for CRISPR/Cas9 gene targeting**

| Target | Left homolgy arm foward | Left homolgy arm reverse | Left homolgy arm sequence | Right homolgy arm foward | Right homolgy arm reverse | Right homolgy arm sequence | gRNA forward | gRNA reverse |
|---|---|---|---|---|---|---|---|---|
| MIER1 N-terminus KI<br><br>Homology arm: TOPO plasmid gRNAs: px459 plasmid | TATATAAAGCTTgactccttggcactgttccttgac | TATATAGCTAGCatttgccgtactgccgggtcacatc | GACTCCTTTGGCACTGTTCCTTGACCTCTCTGATCCGTAGACAACCCCCTCCGCCTGGAGTGGCGGATCAGCTGGAGCCAGCGAAGCGCCCCGCGCGGTTGCCCACCTCCTCCCACACCCACCTTGACTCCGCCCCCTCCGCTCTTCCCGGGGAGGGCTGGCCGCGGGGCCGCGCGCGCCCCTGCTCCGGCGCGTGCTCGCTGGTCTTTTCCCTCCAGTCCAGCCCAGCCGGGGCGCCGCGAGGGGGCGGAGTGGGGTGTGGTGGGCGCGCTCGGGCGGCTCCTGCGCGTTCCCGCCGAGGCAGTGGCGGCGGGAGCGGCAGAGACGGCAGCGGCCGGAGTCCCGTTGCTGAGTCTCACATCCGGGTTCTGGCCGTGACCCAGCTGCGGCCGCCGCGGAGATGTGACCCGGCAGTACGGCAAAT | TATATAACGCGTGAAAACCTGTACTTCCAAAGCatggcggaggtaagggagcgag | TATATATCTAGAacagactggatgtcagtcgcaaatgg | ATGGCGGAGGTAAGGGAGCGAGCTCCCCCTCCCTGTCCCGGAGCCGGGCGCCCCCGGCCCTGGGCCGGGGAGGAGTGGAGTGGGCCTGGCCAGGAGAGGCCCGGCCCTGCTGGCGCCGCTGCCCACCTGTTGTCCGGAACAGGCCATTCTCTGGGCGGGAGCTTCGCCTGGAACTTAGTTGGGATGGGTCCGGGGGTAGGAGGAAGCGGAGGCGTACTTGGACTTCGGGGGCAGTTTGAAGGGTAGTGCAGCGTAGTAGTGCGGTGCGAGGGGGAGGGGGCGCCCGCCGGGGAAGGGGTGCCTGTGGCGCAACTTCGCCGGCCGGTGCTAGGGGCCAGGAGAGAAGCTGCTGCTTCCTCTTGCGAAGTTTTTTCCCTTCCTAACCGAGATGCTGTTGCGCCCTTCCGCGTCACCCCCATCCATTTGCGACTGACATCCAGTCTGT | CACCGCAGTACGGCAAATATGGCGG | aaacCCGCCATATTTGCCGTACTGC |
| KDM1A C-Terminus KI<br><br>Homology arm: TOPO plasmid gRNAs: px330 plasmid | aattaaAAGCTTCTACTTGTCAATTCTGGGAAAGTGCCTAC | TTAATTactagtCATGCTTGGGGACTGCTGTGCAGG | CTACTTGTCAATTCTGGGAAAGTGCCTACTGATAAGGGAGACTCTTCGATAGAATGATGAATAGTAATTGGGGGGGTCAGCCTTTAAAAAGGTCAACAGCAATTTAAGTACTTAGCAATTTAAGTACAAGAATAAAGGTATATGTGCAGCCTGCCAATTTTCTCTTTTTCCCCTAAAATAGCCGATTCCACGACTCTTCTTTGCGGGAGAACATACGATCCGTAACTACCCAGCCACAGTGCATGGTGCTCTGCTGAGTGGGCTGCGAGAAGCGGGAAGAATTGCAGACCAGTTTTTGGGGGCCATGTATACGCTGCCTCGCCAGGCCACACCAGGTGTTCCTGCACAGCAGTCCCCAAGCATG | AATTAAacgcgtGATGCATTCTAAGGGAAGAGGCCC | AATTAATCTAGAGCCCCCATCTTGTATGTAACTCCAAC | GATGCATTCTAAGGGAAGAGGCCCATGTGCCTGTTTCTGCCATGTAAGGAAGGCTCTTCTAGCAATACTAGATCCCACTGAGAAAATCCACCCTGGCATCTGGGCTCCTGATCAGCTGATGGAGCTCCTGATTTGACAAAGGAGCTTGCCTCCTTTGAATGACCTAGAGCACAGGGAGGAACTTGTCCATTAGTTTGGAATTGTGTTCTTCGTAAAGACTGAGGCAAGCAAGTGCTGTGAAATAACATCATCTTAGTCCCTTGGTGTGTGGGGTTTTTGTTTTTTTTTTATATTTTGAGAATAAAACTTCATATAAAATTGGCCCTCTCTTTTGTTCCTTTGAGTTGGAGTTACATACAAGATGGGGGC | CACCGTGAGACAGATGCATTCTAA | aaacTTAGAATGCATCTGTCTCAC |
| DNTTIP1 C-Terminus KI<br><br>Homology arm: TOPO plasmid gRNAs: px330 plasmid | ttaattAAGCTTGATTGTCTCAAGGAACCATTCCCATCC | aattaaTCTAGAGCCCTTGATGTTCTGTCAATTCCCAC | GATTGTCTCAAGGAACCATTCCCATCCCTCCCCACTTACTCTTCCGTACTAAATTGTAAACCGCATCCCCCGCTTCCACCTAGTAGCAGGCCTTACATTAAACAGATGTTTAATGAAGTGTTACTGAGTTGGAGTGAAATGAATCAGGCAATGACCACTCTCCCTTGCTCCTGCCTCAGGCCTACCTCCTCATCGAGGAGGACATCCGGGACCTTGCGGCCAGTGATGATTACAGGTAAAACCAGTGGGTCTCCTGCCCCTTCTCCTCTCCCTGCCTCCAAATAGCCCCTAGAATGAGGCAGGGCCTACATCCTCACTTCCCCCAACTTCTCTCTTCAATGTCTGTTCCAGAGGATGCCTGGATCTGAAGCTAGAGGAATTGAAATCCTTTGTCCTACCCTCCTGGATGGTGGAGAAGATGAGAAAGTATATGGAGACACTACGGACAGAGAATGAGCATCGTGCTGTTGAAGCACCTCCACAGAC | aattaaTCTAGAGCCCTTGATGTTCTGTCAATTCCCAC | ttaattACGCGTGGCCGGGTCCCCTGGCCAC | GGCCGGGTCCCCTGGCCACACTTGGCAGCCCTCCTCCAAAGCCCTCTTCCTCACGTGGCTGAGGCCACCGCTGGGACTGCTCCTAGATGGATCTCAGCGGCATTAAGCTGTGCCTGAGCGAGTTTGTAGTGACTCACTGCACAGCACCCCCAGACTAGCATGTGGTTCTATATTTGTAAAGTTATTGGGATAAGAAACAATTAAACAGTTTGTAGTAAACACAGATGGTGAACCTGCTGTGCCCTCTACCTTGTGGGAATTGACAGAACATCAAGGGC | caccGAAGCACCTCCACAGACCTG | aaacCAGGTCTGTGGAGGTGCTTC |

**Table S1 continued...**

| Target | Forward | Reverse | gBlock_NEO | gBlock_HYG | gRNA forward | gRNA reverse |
|---|---|---|---|---|---|---|
| MTA2 C-Terminus KI<br><br>Homology arm: TOPO plasmid gRNAs: px330 plasmid | aattaaAAGCTTTGGGCTTCACCTGGAGGTCTTAA | TTAATTTCTAGAATGGGCCAAGTTCCTGCAGTCTGT | TGGGCTTCACCTGGAGGTCTTAATGTTCTCTAAGCCTTTTCTGTTTCCCACTTGCTCTTTCTGGCCAGGGCCCTACGGAAGGCTCTGACCCATCTGGAAATGCGGCGAGCTGCTCGCCGACCCAACTTGCCCCTGAAGGTGAAGCCAACGCTGATTGCAGTGCGGCCCCCTGTCCCTCTACCTGCACCCTCACATCCTACCAGCACCAATGAGCCTATTGTCCTGGAGGACGGTGGCGGAGGGTCTGGTGGCGGAGGGTCTggaagcggaGAAAACCTGTACTTCCAAAGCGGTGGCGGAGGTTCAggagtgcaggtggaaaccatctccccaggagacgggcgcaccttccccaagcgcggccagacctgcgtggtgcactacaccgggatgcttgaagatggaaagaaagttgattcctcccgggacagaaacaagccctttaagtttatgctaggcaagcaggaggtgatccgaggctgggaagaaggggttgcccagatgagtgtgggtcagagagccaaactgactatatctccagattatgcctatggtgccactgggcacccaggcatcatcccaccacatgccactctcgtcttcgatgtggagcttctaaaactggaaggaagcggaGACTACAAAGATCACGACGGAGACTATAAAGATCACGACATCGATTATAAAGATGACGACGATAAAGGATCAGGGGCCACTAACTTTTCCCTGCTGAAGCAGGCCGGAGACGTGGAGGAGAACCCCGGGCCCGCTAGCATGAAAAAGCCTGAACTCACCgcgacgtctgtcgagaagtttcTGATCGAAAAGTTCGACAGCGTCTCCGACCTGATGCAGCTCTCGGAGGGCGAAGAATCTCGTGCTTTCAGCTTCGATGTAGGAGGGCGTGGATATGTCCTGCGGGTAAATAGCTGCGCCGATGGTTTCTACAAAGATCGTTATGTTTATCGGCACTTTGCATCGGCCGCGCTCCCGATTCCGGAAGTGCTTGACATTGGGGAGTTCAGCGAGAGCCTGACCTATTGCATCTCCCGCCGTGCACAGGGTGTCACGTTGCAAGACCTGCCTGAAACCGAACTGCCCGCTGTTCTTCAGCCGGTCGCGGAGGCTATGGATGCGATCGCTGCGGCCGATCTTAGCCAGACGAGCGGGTTCGGCCCATTCGGACCGCAAGGAATCGGTCAATACACTACATGGCGTGATTTCATATGCGCGATTGCTGATCCCCATGTGTATCACTGGCAAACTGTGATGGACGACACCGTCAGTGCGTCCGTCGCGCAGGCTCTCGATGAGCTGATGCTTTGGGCCGAGGACTGCCCCGAAGTCCGGCACCTCGTGCACGCGGATTTCGGCTCCAACAATGTCCTGACGGACAATGGCCGCATAACAGCGGTCATTGACTGGAGCGAGGCGATGTTCGGGGATTCCCAATACGAGGTCGCCAACATCTTCTTCTGGAGGCCGTGGTTGGCTTGTATGGAGCAGCAGACGCGCTACTTCGAGCGGAGGCATCCGGAGCTTGCAGGATCGCCACGCCTCCGGGCGTATATGCTCCGCATTGGTCTTGACCAACTCTATCAGAGCTTGGTTGACGGCAATTTCGATGATGCAGCTTGGGCGCAGGGTCGATGCGACGCAATCGTCCGATCCGGAGCCGGGACTGTCGGGCGTACACAAATCGCCCGCAGAAGCGCGGCCGTCTGGACCGATGGCTGTGTAGAAGTACTCGCCGATAGTGGAAACCGACGCCCCAGCACTCGTccgagggcaaaggaaTGAACGCGTGCACCTGTGGGGAAGGGAGGTGGGCTGAGAGGTAGAGGGTGGATGCCCAGGGCACCCAAACCTCCCTTCCCTTTCGTGTCGAAGGGAGTGAGGAGTGAATTAAGGAAGAGAGCAAGTGAGTGTGTGTCCCTGGAGGGGTTGGGCGCCCTCTGGTGTTACCACCTCGAGACTTGTCTCATGCCTCCATGCTTGCCGATGGAGGACAGACTGCAGGAACTTGGCCCAT | TGGGCTTCACCTGGAGGTCTTAATGTTCTCTAAGCCTTTTCTGTTTCCCACTTGCTCTTTCTGGCCAGGGCCCTACGGAAGGCTCTGACCCATCTGGAAATGCGGCGAGCTGCTCGCCGACCCAACTTGCCCCTGAAGGTGAAGCCAACGCTGATTGCAGTGCGGCCCCCTGTCCCTCTACCTGCACCCTCACATCCTACCAGCACCAATGAGCCTATTGTCCTGGAGGACGGTGGCGGAGGGTCTGGTGGCGGAGGGTCTggaagcggaGAAAACCTGTACTTCCAAAGCGGTGGCGGAGGTTCAggagtgcaggtggaaaccatctccccaggagacgggcgcaccttccccaagcgcggccagacctgcgtggtgcactacaccgggatgcttgaagatggaaagaaagttgattcctcccgggacagaaacaagccctttaagtttatgctaggcaagcaggaggtgatccgaggctgggaagaaggggttgcccagatgagtgtgggtcagagagccaaactgactatatctccagattatgcctatggtgccactgggcacccaggcatcatcccaccacatgccactctcgtcttcgatgtggagcttctaaaactggaaggaagcggaGACTACAAAGATCACGACGGAGACTATAAAGATCACGACATCGATTATAAAGATGACGACGATAAAGGATCAGGGGCCACTAACTTTTCCCTGCTGAAGCAGGCCGGAGACGTGGAGGAGAACCCCGGGCCCGCTAGCATGCCTGTAATTTCTACCCAGACTGGACGGgccatgattgagcaagacgggctCCACGCTGGCAGCCCCGCAGCTTGGGTCGAGCGACTGTTCGGGTACGATTGGGCACAGCAGACAATAGGGTGCAGCGATGCCGCCGTCTTCCGGCTCAGCGCGCAAGGCCGGCCTGTCCTGTTTGTTAAAACCGATCTGAGCGGGGCCCTGAACGAACTGCAGGATGAGGCGGCTAGACTTAGCTGGCTTGCGACCACCGGAGTGCCGTGTGCTGCCGTTCTGGACGTCGTAACAGAGGCGGGAAGGGATTGGCTGCTGCTCGGGGAGGTCCCTGGCCAAGATTTGTTGTCCTCCCACCTGGCACCTGCAGAGAAGGTAAGCATCATGGCAGATGCCATGCGCAGGCTGCACACCCTGGATCCCGCCACGTGTCCTTTCGACCACCAGGCCAAGCACCGAATTGAGAGGGCCAGGACACGCATGGAGGCCGGCCTGGTGGATCAGGACGATCTTGACGAGGAACATCAGGGCCTCGCCCCAGCGGAGCTCTTTGCTCGGCTGAAAGCTAGAATGCCTGATGGTGAAGATCTCGTCGTGACCCACGGAGATGCCTGCCTGCCCAACATCATGGTAGAAAACGGACGCTTCTCTGGCTTTATCGATTGTGGCCGGCTTGGAGTTGCTGATAGATATCAGGACATTGCACTCGCGACAAGAGACATTGCCGAGGAACTCGGTGGTGAATGGGCAGACCGGTTCCTGGTGCTGTACGGGATCGCTGCCCCTGACTCACAGAGGATCGCATTTTACAggttgctggacgaattttttaaACGCGTGCACCTGTGGGGAAGGGAGGTGGGCTGAGAGGTAGAGGGTGGATGCCCAGGGCACCCAAACCTCCCTTCCCTTTCGTGTCGAAGGGAGTGAGGAGTGAATTAAGGAAGAGAGCAAGTGAGTGTGTGTCCCTGGAGGGGTTGGGCGCCCTCTGGTGTTACCACCTCGAGACTTGTCTCATGCCTCCATGCTTGCCGATGGAGGACAGACTGCAGGAACTTGGCCCAT | CACCGTGGAGGACTGAGCACCTGTG | aaacCACAGGTGCTCAGTCCTCCAC |

| Target | Forward | Reverse (Blasticidin) | Reverse (Puromycin) | gRNA forward | gRNA reverse |
|---|---|---|---|---|---|
| HDAC3 C-Terminus KI<br><br>Cloning in Pitch plasmids gRNAs: px330 plasmid | ccgcgttacatagcatcgtacgcgtacgtgtttggGACAATGACAAGGAAAGCGATGTGGAGATTggtggcggtggctcgggcgg | cagcattctagagcatcgtacgcgtacgtgtttggAGAAATTCCTTGGGACACAGCATCCCAAGCtcagccctcccacacataac | cagcattctagagcatcgtacgcgtacgtgtttggAGAAATTCCTTGGGACACAGCATCCCAAGCtcaggcaccgggcttgcggg | CACCGTGGAGATTTAAGAGTGGCT | AAACAGCCACTCTTAAATCTCCAC |

| Target | Forward (Blasticidin) | Forward (Puromycin) | Reverse | gRNA forward | gRNA reverse |
|---|---|---|---|---|---|
| Sin3A C-Terminus KI<br><br>Cloning in Pitch plasmids gRNAs: px330 plasmid | TTACATAGCATCGTACGCGTACGTGTTTGGTCCCTGAGCATGAGCACAGAATGAAGCGGCGAATGCCAAGCCTTTGTCTCAAGAAGAATCCACCCTCATTGAAAGAGCAACGG | TTACATAGCATCGTACGCGTACGTGTTTGGTCCCTGAGCATGAGCACAGAATGAAGCGGCGAATGACCGAGTACAAGCCCACGGTGCGCCTCGCCACCC | TTCTAGAGCATCGTACGCGTACGTGTTTGGCATACACCGGTGACTCCTGGTCATCCAAACGAGATCCGCCGCCACCCGACCCACCACCGCCCGAGCCACCGCCACCTTCCAG | CACCGCACAGAATGAAGCGGCGTT | AAACAACGCCGCTTCATTCTGTGC |
| MIER2 KO<br><br>Cloning in Pitch plasmids gRNAs: px459 plasmid | | | | CACCGCCGGGCTTGCAGACAACAGC | aaacGCTGTTGTCTGCAAGCCCGGC |
| MIER3 KO<br><br>Cloning in Pitch plasmids gRNAs: px459 plasmid | | | | CACCGAAGAGGAAATGATGGATGA | aaacTCATCCATCATTTCCTCTTC |

**Table S1 continued…**

| Target | Forward (Blasticidin) | Forward (Puromycin) | Reverse | gRNA forward | gRNA reverse |
|---|---|---|---|---|---|
| MTA1 KO | | | | | |
| Cloning in Pitch plasmids gRNAs: px459 plasmid | | | | CACCGATACCTGATCCGGAGAATCG | aaacCGATTCTCCGGATCAGGTATC |
| MTA3 KO | | | | | |
| Cloning in Pitch plasmids gRNAs: px330 plasmid | | | | CACCGTTCTTACCTGATATGTGTTG | aaacCAACACATATCAGGTAAGAAC |

**Table S2. Antibodies used in the study**

| Target | Host/Isotype | Clonality | Experiment | Concentration | Source | Catalogue # |
|---|---|---|---|---|---|---|
| FLAG | Mouse/IgG | Monoclonal | Western Blotting | 1:1000 | Merck | F1804 |
| SIN3A | Rabbit/IgG | Monoclonal | Western Blotting | 1:1000 | Abcam | AB129087 |
| MTA3 | Rabbit/IgG | Monoclonal | Western Blotting | 1:1000 | Abcam | Ab176346 |
| MTA2 | Mouse/IgG | Monoclonal | Western Blotting | 1:1000 | Santa Cruz | SC-55599 |
| MTA1 | Rabbit/IgG | Monoclonal | Western Blotting | 1:1000 | Cell Signalling | 5647S |
| LSD1 | Rabbit/IgG | Monoclonal | Western Blotting | 1:1000 | Abcam | Ab129195 |
| DNTTIP1 | Rabbit/IgG | Polyclonal | Western Blotting | 1:1000 | Abcam | Ab174663 |
| MIER1 | Rabbit/IgG | Polyclonal | Western Blotting | 1:1000 | Invitrogen | PA5-40896 |
| MIER3 | Rabbit/IgG | Polyclonal | Western Blotting | 1:1000 | Invitrogen | PA5-31879 |
| HDAC3 | Rabbit/IgG | Monoclonal | Western Blotting | 1:1000 | Abcam | ab76295 |
| TBP | Mouse/IgG | Monoclonal | Western Blotting | 1:300 | Santa Cruz | SC-421 |
| RCoR1 | Mouse/IgG | Monoclonal | Western Blotting | 1:1000 | Merck | MABN486 |
| HDAC1 | Rabbit/IgG | Monoclonal | Western Blotting | 1:1000 | Abcam | Ab109411 |
| HDAC2 | Mouse/IgG | Monoclonal | Western Blotting | 1:1000 | Merck | 05-814 |
| MIDEAS | Rabbit/IgG | Polyclonal | Western Blotting | 1:500 | Merck | HPA003111 |
| LSD1 | Rabbit/IgG | Monoclonal | co-IP | 2ug | Abcam | Ab129195 |
| FLAG | Mouse/IgG | Monoclonal | co-IP | 10ug | Merck | F1804 |
